# Cdc42EP5/BORG3 modulates SEPT9 to promote actomyosin function and melanoma invasion and metastasis

**DOI:** 10.1101/570747

**Authors:** Aaron J Farrugia, Javier Rodríguez, Jose L Orgaz, María Lucas, Victoria Sanz-Moreno, Fernando Calvo

## Abstract

Fast amoeboid migration in the invasive fronts of melanoma is controlled by high levels of actomyosin contractility, which underlie its highly metastatic potential. How this migratory behaviour is coupled to other cytoskeletal components is poorly understood. Septins are increasingly recognized as novel cytoskeletal components, but details on their regulation and contribution to cancer migration and metastasis are lacking. Here, we show that the septin regulator Cdc42EP5 is consistently required for melanoma cells to migrate and invade into collagen-rich matrices, and to locally invade and disseminate *in vivo*. Cdc42EP5 associates with actin structures leading to increased actomyosin contractility and amoeboid migration. Cdc42EP5 effects these functions through SEPT9-dependent F-actin crosslinking, which enables the generation of F-actin bundles required for the sustained stabilisation of highly contractile actomyosin structures. This study provides evidence for Cdc42EP5 as a regulator of cancer cell motility that coordinates actin and septin networks. It also describes a unique role for SEPT9 in invasion and metastasis, and illustrates a mechanism that regulates its function in melanoma.

## Introduction

Malignant melanoma is a very aggressive type of skin cancer due to its highly metastatic behaviour, which relies in the increased ability of melanoma cells to migrate and invade (1). Melanoma can migrate as single cells where they display two major morphologies: elongated-mesenchymal or rounded-amoeboid. Mesenchymal migration is characterised by Rac-driven actin-based protrusions, matrix degradation and strong focal adhesions (FAs) coupled to actin fibres that enable force transmission (2). In contrast, amoeboid migration modes are characterised by a rounded morphology as well as blebs, lower levels of adhesion, and high levels of actomyosin contractility. Importantly, amoeboid behaviour is prominent in the invasive fronts of melanomas in animal models (3–5) and human lesions (5–7). It has also been associated with increased risk of metastasis and poorer prognosis (7), which underlies the need for a better mechanistic understanding of the process. Actomyosin contractility driven by the motor protein Myosin II is critical for rounded migration (8). This process has been shown to be tightly controlled by Rho-ROCK signalling leading to increased phosphorylation of the regulatory myosin light chain (MLC2)(9). However, how actin structures are organized and coordinated with other cytoskeletal components to enable their correct assembly and the formation of fully functional actomyosin networks is not well understood.

Septins are a large conserved family of GTP binding proteins that participate in a broad spectrum of cellular functions (10). Septins have been proposed as the fourth component of the cytoskeleton due to their ability to form higher-order structures such as filaments, which can associate with distinct subsets of actin filaments and microtubules, as well as membranes of specific curvature and composition (11). Importantly, septins are emerging as crucial regulators of the generation, maintenance and positioning of cytoskeletal networks with potential roles in cell migration. In line with this, different septins have been shown to be required for mesenchymal migration in epithelial and endothelial cells (12, 13). In addition, septins form a uniform network at the cell cortex in leukocytes and SEPT7 expression is required for rapid cortical contraction during dynamic shape changes (14, 15). In cancer, a potential role for septins in modulating aggressiveness is also starting to emerge (16, 17), although little is known about the molecular details and functions of individual members in melanoma.

Although a role for septins in modulating cytoskeletal rearranges is clearly emerging, the regulatory mechanisms required for establishing and maintaining these interactions in migratory cells are still elusive. Binder-of-Rho-GTPases (Borg) proteins (also called Cdc42 effector proteins or Cdc42EPs) are amongst the few proteins known to interact with septins and regulate their function (18). Borg proteins vary in length, but all contain Borg Homology 3 Domain (BD3) that binds septins (18). Although they remain largely uncharacterised, recent studies suggest crucial roles of Borg proteins in regulating cytoskeletal organization and related cellular processes (13, 19). Yet, the exact role of Borg proteins and septins in cancer, and their relationship in modulating cancer cell invasion and metastatic dissemination remains to be determined.

Here, we investigate the contribution of individual Borg proteins to rounded-amoeboid cell migration and invasion using melanoma models. We find that Cdc42EP5 is consistently required for these processes *in vitro*, and for local invasion and metastasis *in vivo*. Mechanistically, we demonstrate that Cdc42EP5 acts by inducing actin cytoskeleton rearrangements that potentiate actomyosin function in a septin-dependent manner, and we show a unique role for SEPT9 in controlling actomyosin contractility and invasion in melanoma.

## Results

### Identification of Cdc42ep5 as a regulator of melanoma migration, invasion and metastasis

To assess the relevance of Borg proteins Cdc42ep1-5 in cancer cell migration and invasion, we first analysed transwell migration after RNAi silencing of individual genes in the murine melanoma model 690.cl2 (20, 21). Note that when referring to murine RNA/proteins we used an uppercase letter followed by all lowercase letters (i.e. Cdc42EP5 or abbreviated forms, Ep5); when referring to human products or in general we used uppercase letters (i.e. CDC42EP5 or EP5). We achieved knockdown of all Borg genes when each gene was specifically targeted (**Fig EV1A**). **Fig 1A** shows that knocking-down Cdc42ep3 and Cdc42ep5 significantly reduced the ability of 690.cl2 cells to migrate through transwell pores, whereas silencing Cdc42ep2 increased migration. To assess how these defects were affecting the ability of melanoma cells to invade, we employed collagen invasion assays and observed that only Cdc42ep5 depletion significantly reduced 690.cl2 invasion (**Fig 1B**). Silencing Cdc42ep5 expression with two independent RNAi in 690.cl2 cells (**Fig 1C; Fig EV1B**) yielded similar results (**Fig 1D and E**). Using CRISPR/CAS9 technology (**Fig EV1C and D**), we generated *Cdc42ep5* knock-out 690.cl2 cells expressing GFP (690.cl2^KO^-GFP) and knock-out cells ectopically re-expressing an N-terminal GFP-Cdc42ep5 fusion (690.cl2^KO^-GFP-Cdc42ep5). Critically, GFP-Cdc42ep5 expression in the reconstituted 690.cl2^KO^ cells was similar to endogenous Cdc42ep5 levels in parental 690.cl2 cells. Functional characterization informed that Cdc42ep5 depletion affected melanoma migration *in vitro* and that these defects were abrogated when Cdc42ep5 expression was reconstituted (**Fig EV1E**). There was also a significant increase in the migratory abilities of parental 690.cl2 cells after Cdc42ep5 over-expression (**Fig EV1F**), confirming a role for this protein in melanoma migration and invasion *in vitro*. These results were further validated in an alternative human melanoma model (WM266.4) and in a model of breast cancer (MDA-MB-231), where *CDC42EP5* was consistently required for migration and invasion (**Fig EV1G-I**).

**Figure 1.**
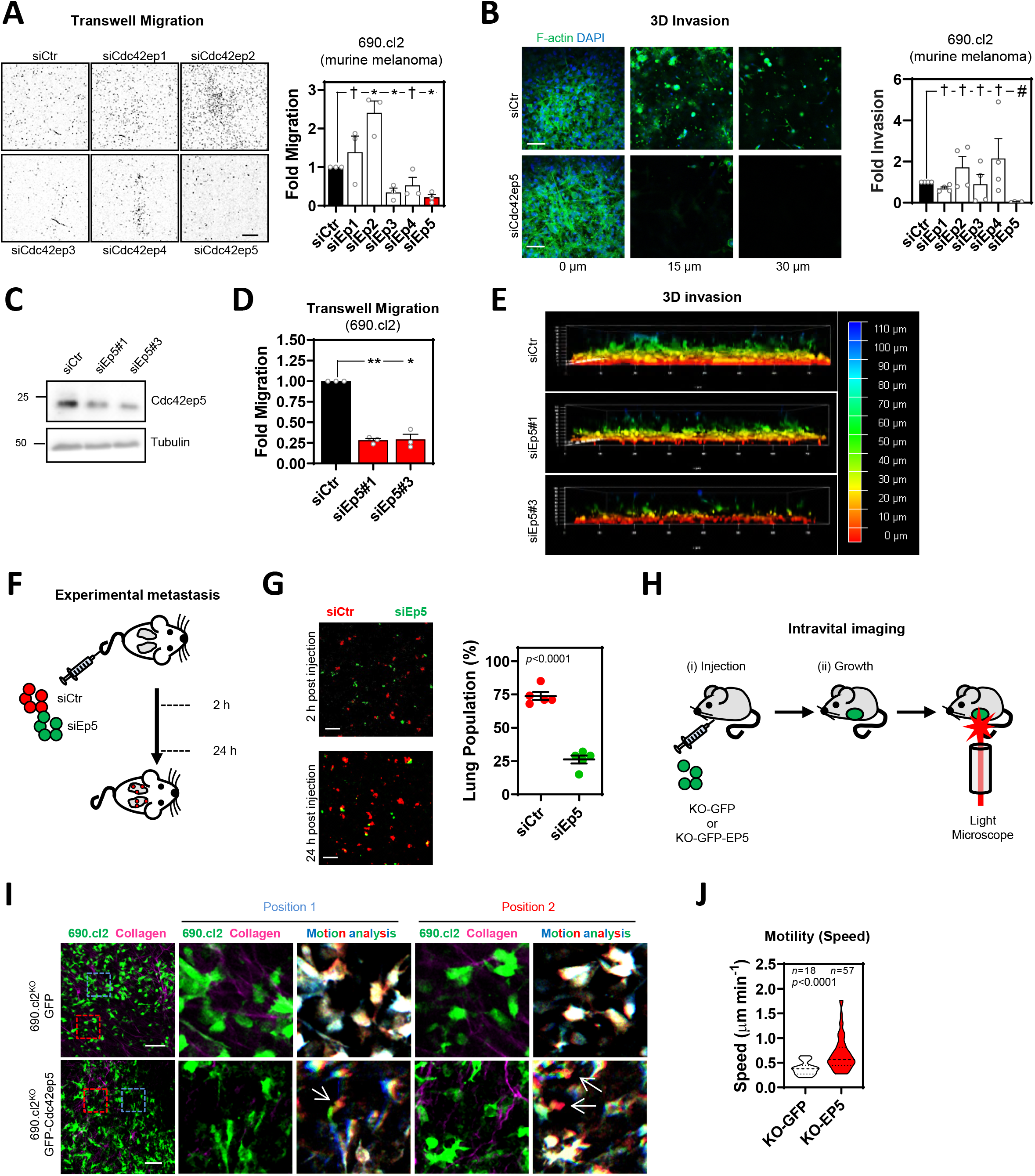
Directed siRNA screen identifies Cdc42EP5 as a new regulator of melanoma migration and invasion. **(A)** Images show DAPI staining of migrated 690.cl2 murine melanoma after transfection with control (siCtr) or Cdc42ep1-5 (siEp1-5) siRNA. Scale bar, 1 mm. Graph shows fold migration. Bars indicate mean ± SEM (*n*=3 independent experiments; one-way paired ANOVA, Dunnet’s test: †, not significant; *, *p* < 0.05). **(B)** Images show F-actin (green) and DAPI (blue) staining of 690.cl2 cells at different levels of collagen invasion after transfection with indicated siRNAs. Scale bar, 100 μm. Graph shows fold invasion. Bars indicate mean ± SEM (*n*=3 independent experiments; one-way paired ANOVA, Dunnet’s test: †, not significant; #, *p* < 0.0001). **(C)** Representative Western blot showing indicated protein levels in 690.cl2 cells after transfection with control (siCtr) or two individual Cdc42ep5 (siEp5) siRNA. **(D)** Fold migration of 690.cl2 cells after transfection with indicated siRNAs. Bars indicate mean ± SEM (*n*=3 independent experiments; one-way paired ANOVA, Dunnet’s test: *, *p* < 0.05; **, *p* < 0.01). **(E)** Images show 3D reconstructions of collagen invasion assays of 690.cl2 cells transfected with indicated siRNAs. Colour scale bar indicates depth of invasion. **(F)** Experimental metastasis approach: equal numbers of control (siCtr, *red*) and Cdc42ep5-knock-down (siEp5, *green*) 690.cl2 cells were injected in the tail vein of mice. 2 h and 24 h after injection, lungs were analysed by microscopy. **(G)** Images of mouse lungs at 2 and 24 h after tail injection with 690.cl2 cells as in (F). Scale bars, 100 μm. Graph shows proportions (%) of siCtr and siEp5 cells in lungs at 24 h. Lines represent mean ± SEM (*n*=5 experiments; unpaired *t* test). **(H)** Intravital imaging approach: 690.cl2^KO^-GFP or 690.cl2^KO^-GFP-Cdc42ep5 cells were injected subcutaneously in mice. Grown living tumours on anaesthetised mice were then exposed and imaged using 2-photon microscopy. **(I)** Intravital imaging of 690.cl2^KO^-GFP or 690.cl2^KO^-GFP-Cdc42ep5 tumours (green, melanoma cells; magenta, collagen fibres). Right panels show motion analysis images from different time points where static cells appear white; arrows indicate fast moving cells represented as a trail of blue-green-red pseudocolours. Scale bar, 100 μm. **(J)** Violin plot showing individual speed of moving cells from (I); *n*, moving cells in 5 mice per condition; Mann-Whitney test).

Melanoma cells can invade locally and disseminate to distant organs including the lungs (22). To confirm the requirement for Cdc42ep5 in these processes *in vivo*, we first investigated lung metastasis after tail vein injection of control and Cdc42ep5-knock-down 690.cl2 cells in mice (**Fig 1F**). Two hours after injection, equal numbers of control and Cdc42ep5-depleted cells lodged in the lungs (**Fig 1G**). However, significantly fewer Cdc42ep5 knock-down cells were present in the lungs after 24 h when compared to control cells, confirming a requirement for Cdc42ep5 in metastasis. Next, we investigated melanoma motility and local invasion using intravital imaging in living tumours (**Fig 1H**). For this long term assay, we employed the CRISPR-engineered cell lines 690.cl2^KO^-GFP and 690.cl2^KO^-GFP-Cdc42ep5, which will allow for assessing the specific role of Cdc42ep5 and discard clonal effects derived from knock-out generation. No changes in melanoma proliferation or tumour growth were observed after modulating Cdc42ep5 expression (**Fig EV1J and K**). Intravital imaging showed an increase in the speed of moving cells in 690.cl2^KO^-GFP-Cdc42ep5 cells when compared to 690.cl2^KO^-GFP cells (**Fig 1I and J, Movie EV1&2**). Together, these results indicate that Cdc42ep5 is consistently required for melanoma migration and invasion *in vitro* and for local invasion and metastatic dissemination *in vivo*.

### Cdc42ep5 promotes actomyosin function in collagen-rich matrices

Fast motility of melanoma cells *in vivo* has been associated with a particular rounded-amoeboid behaviour that promotes local invasion and metastasis, negatively impacting patient survival (4, 23). In melanoma, amoeboid migration *in vivo* is tightly regulated by actomyosin function (3–5). Importantly, rounded-amoeboid behaviour in melanoma can be modelled *in vitro* by seeding cells on collagen-rich matrices and assessing cell morphology and actomyosin function (4, 24).

To explore whether Cdc42ep5 was promoting migration and invasion by participating in these processes, we first used confocal microscopy to determine the precise localization of Cdc42ep5 with respect to actomyosin networks in 690.cl2 cells seeded on collagen-rich matrices. We could not identify suitable Cdc42ep5 antibodies for immunofluorescence, so we stably expressed our GFP-Cdc42ep5 construct in 690.cl2 cells to investigate its cellular localization. Initial analyses informed that individual cells presented different degrees of Cdc42ep5 expression, which were positively correlated with actomyosin function (as measured by pS19-MLC2 intensity levels), F-actin levels and cell roundness, suggesting a potential role for Cdc42ep5 in regulating these parameters (**Fig EV2A and B**). In depth confocal analysis of Cdc42ep5-expressing rounded 690.cl2 cells on collagen-rich matrices informed that GFP-Cdc42ep5 localized preferentially at the cell cortex (**Fig 2A**), co-localizing with two key components of cortical actomyosin networks, F-actin and pS19-MLC2. Actomyosin contractility at the cellular cortex drives the formation of protrusions known as blebs (25). These blebs have been implicated in enhanced migration thorough complex environments, and have been observed in highly invasive melanoma cells (4). Using high resolution confocal microscopy, GFP-Cdc42ep5 was found to localize with F-actin on the cortex at areas of blebbing activity within the cell (**Fig EV2C**).

**Figure 2.**
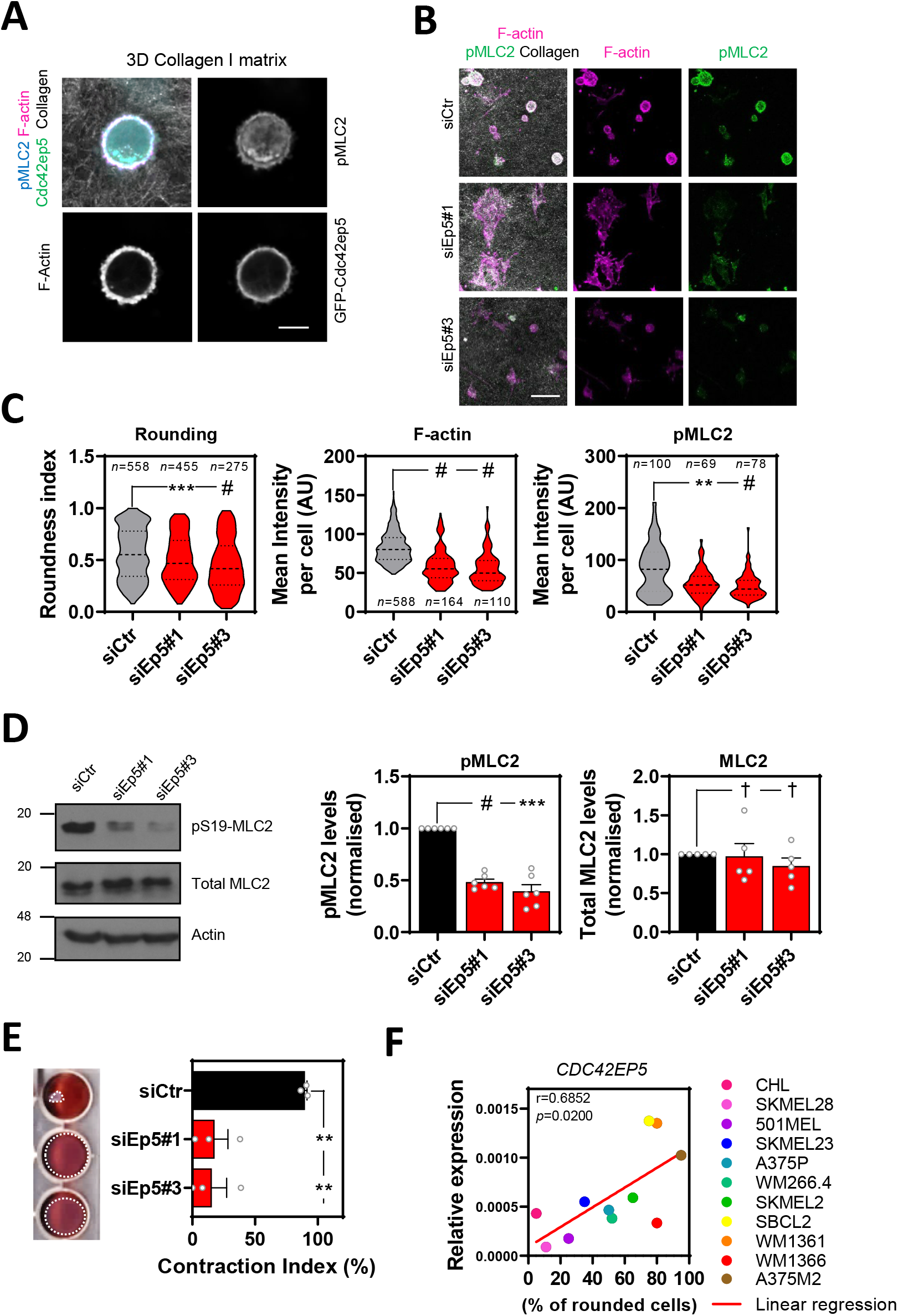
Cdc42EP5 modulates actomyosin function in melanoma in 3D. **(A)** Image of a 690.cl2 cell on collagen-rich matrix expressing GFP-Cdc42ep5 (green) and stained for F-actin (magenta) and pS19-MLC2 (cyan); collagen fibres (grey) were captured by reflectance signal. Individual greyscale channels are also shown. Scale bar, 10 μm. **(B)** Images of 690.cl2 cells on collagen-rich matrices after transfection with control (siCtr) or two individual Cdc42ep5 (siEp5). Images show staining for F-actin (magenta) and pS19-MLC2 (green); collagen fibres (grey) were captured by reflectance signal. Scale bar, 40 μm. **(C)** Violin plots showing roundness index (*left*), F-actin mean intensity (*middle*) and pS19-MLC2 mean intensity (*right*) of individual cells from (B); *n* represents individual cells; Kruskal-Wallis and Dunn’s tests: **, *p* < 0.01; ***, *p* < 0.001; #, *p* < 0.0001. **(D)** Western blots showing indicated protein levels in 690.cl2 cells after transfection with control (siCtr) and two individual Cdc42ep5 (siEp5) siRNAs. Graphs show quantification of normalized pS19-MLC2 and total MLC2 levels. Bars indicate mean ± SEM (*n*=6 [pMLC2] and 5 [MLC2] independent experiments; one-way paired ANOVA, Dunnet’s test: †, not significant; ***, *p* < 0.001; #, *p* < 0.0001). **(E)** Images and graph showing gel contraction of 690.cl2 cells after indicated treatments. Bars indicate mean ± SEM (*n*=3 independent experiments; one-way paired ANOVA, Dunnet’s test: **, *p* < 0.01). **(F)** Graph shows expression of *CDC42EP5* mRNA normalized in human melanoma cell lines with increasing rounding coefficients. Person correlation coefficient (*r*), statistical significance (*p*) and linear regression (red line) are shown. Each point in the graph represents the mean value of 3 independent experiments.

A role for Cdc42ep5 in modulating actomyosin contractility in 3D was further confirmed by perturbation analyses. Thus, silencing Cdc42ep5 with two independent RNAi significantly decreased the roundness index of 690.cl2 cells seeded on collagen-rich matrices (**Fig 2B and C; Fig EV2D and E**). Importantly, the reduction in cell rounding after Cdc42ep5 depletion was associated with a decrease in F-actin and pS19-MLC2 levels by immunofluorescence (**Fig 2B and C**). Immunoblot analyses of whole cell lysates confirmed the reduction of pS19-MLC2 but not total MLC2 levels after Cdc42ep5 silencing (**Fig 2D**). As a result, the ability of cells to contract collagen-rich matrices was severely impeded after Cdc42ep5 depletion (**Fig 2E**), which underlines the relevance of Cdc42ep5 in modulating actomyosin activity and cellular contractility. The association between cell rounding and Cdc42EP5 expression was consistent in other melanoma cell lines. Thus, we found that *CDC42EP5* was the only Borg gene whose expression was significantly correlated with cell roundness (as assessed on cells seeded on collagen-rich matrices) in a panel of 11 human melanoma cell lines of varying degrees of rounding (6)(**Fig 2F; Fig EV2F**). Perturbation analyses confirmed this observation, as silencing Borg genes *CDC42EP1-4* in 690.cl2 had no effect on pS19-MLC2 levels, which contrasted with the significant decrease after *CDC42EP5* depletion (**Fig EV2G**). Together, these data indicated that CDC42EP5 promotes the generation of an actomyosin cortex, which impacts actomyosin activity and cell morphology, and induces invasive phenotypes in melanoma cells in 3D settings.

### Cdc42ep5 regulates the organization of actomyosin networks and is required for the maturation of FAs

Whilst actomyosin contractility in 3D pliable environments drives rounded-amoeboid behaviour, in stiff undeformable 2D culture it can be associated with the formation of stress fibres, long actin filaments decorated with active MLC2 (pS19-MLC2)(26). To assess if actomyosin was regulated by Cdc42ep5 independently of the physical context, we next used 2D culture. We observed that Cdc42ep5 formed an intricate filamentous network that overlapped fibrillary actin bundles in basal the perinuclear region in 690.cl2 cells (**Fig 3A**). Cdc42ep5 filaments were less evident towards the cell periphery. Time-lapse imaging showed that Cdc42ep5 filaments were dynamic and aligned with F-actin fibres, stretching along the cell body towards the cell periphery (**Movie EV3**). Cdc42ep5 filaments were absent from lamellipodial regions and membrane ruffles. In agreement, Pearson’s correlation analyses informed that Cdc42ep5 co-localized with F-actin preferentially in the perinuclear region in fixed cells (**Fig 3A and B**). There was no co-localization of GFP-Cdc42ep5 with acetylated alpha-tubulin, therefore direct associations between Cdc42EP5 and microtubules were unlikely (**Fig EV3A; Fig 3B**). More detailed analyses of cellular structures in 2D informed that GFP-Cdc42ep5 filaments aligned with both F-actin- and pS19-MLC2-positive filaments (i.e. stress fibres)(**Fig EV3B**). In cellular protrusions, Cdc42ep5 localized with actomyosin filaments mostly around the edges of the protrusion. Confocal analysis of the perinuclear region indicated that Cdc42ep5 filaments stretched across the basal side in alignment with F-actin fibres, with activated MLC2 dotted around the network. Cdc42ep5 also localized in the apical region, where it formed a ‘wavy’ network around actomyosin filaments.

**Figure 3.**
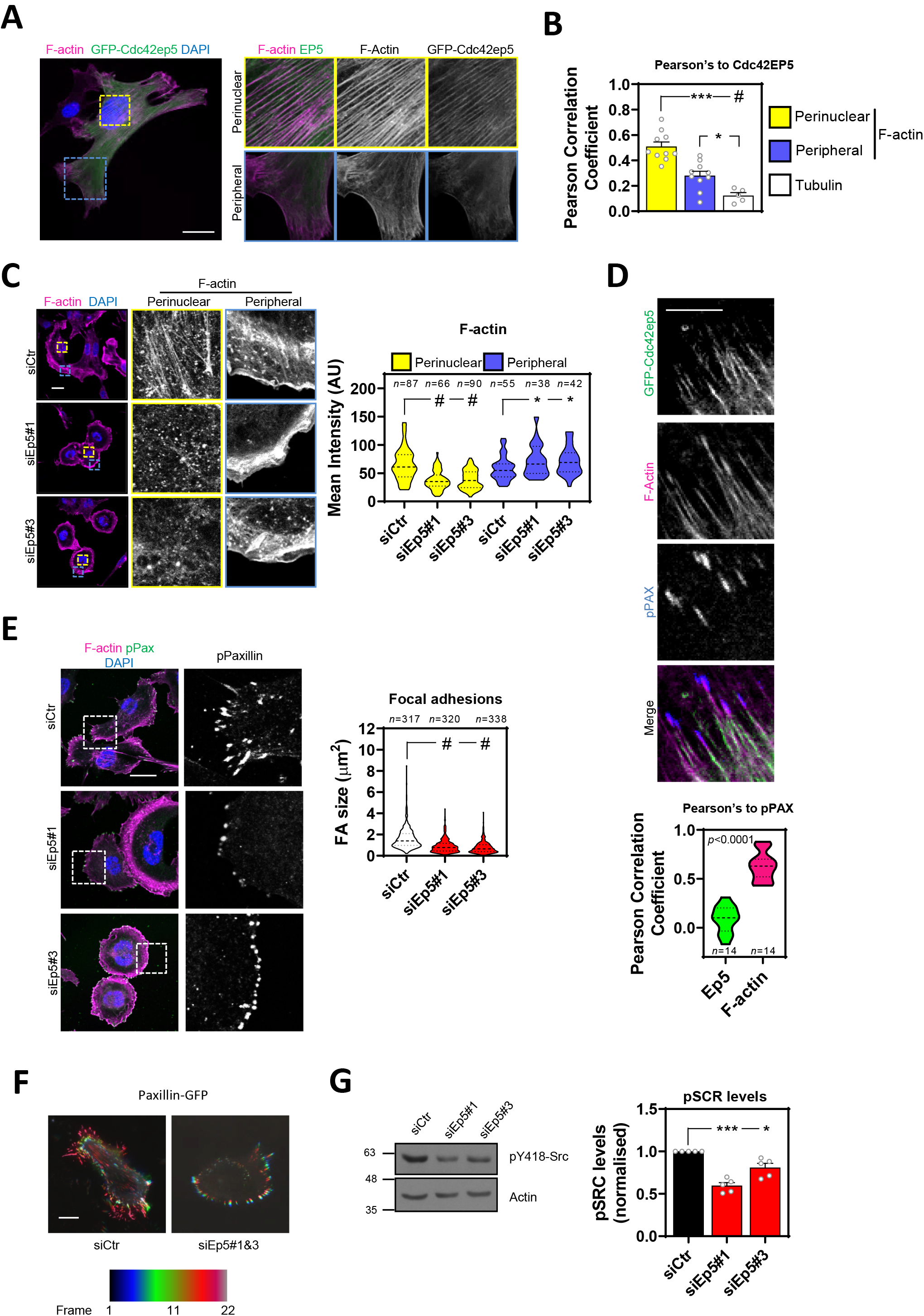
Cdc42EP5 promotes stress fibre formation and focal adhesion maturation in 2D. **(A)** Image shows a 690.cl2 cell expressing GFP-Cdc42EP5 (green) on glass and stained for F-actin (magenta) and DAPI (blue). Additional panels show magnifications of perinuclear and peripheral areas. Scale bar, 25 μm. **(B)** Graph shows Pearson Correlation coefficient of GFP-Cdc42ep5 signal against perinuclear and peripheral F-actin staining, and tubulin staining. Bars indicate mean ± SEM (*n*=10 independent cells [perinuclear and peripheral F-actin] and 5 [tubulin]; one-way paired ANOVA, Dunnet’s test: *, *p* < 0.05; ***, *p* < 0.001; #, *p* < 0.0001). **(C)** Images of 690.cl2 cells after transfection with control (siCtr) and two individual Cdc42ep5 (siEp5) siRNAs seeded on glass. Images show F-actin (magenta) and DAPI (blue) staining with zoom-up panels showing perinuclear and peripheral regions. Scale bar, 25 μm. Graph is a violin plot quantification of F-actin intensity in the perinuclear and peripheral regions in the indicated points (*n*, specific regions in individual cells; Kruskal-Wallis and Dunn’s tests: *, *p* < 0.05; #, *p* < 0.0001). **(D)** Images show individual and merged channels of 690.cl2 expressing GFP-Cdc42ep5 (green) on glass and stained for F-actin (magenta) and pY118-Paxilin (blue). Scale bar, 5 μm. Graph at the bottom is a violin plot showing the Pearson Correlation coefficient of Cdc42ep5 and F-actin staining against pY118-Paxilin-positive areas (*n*, pY118-Paxilin-positive areas in individual cells; unpaired *t* test). **(E)** Images of 690.cl2 cells on glass. Images show F-actin (magenta), pY118-Paxilin (green) and DAPI (blue) staining. Single channel magnification of pY118-Paxilin staining is also shown. Scale bar, 25 μm. Graph is a violin blot showing quantification of focal adhesion (FA) size i.e. pY118-Paxilin positive area (*n*, peripheral regions in individual cells; Kruskal-Wallis and Dunn’s: #, *p* < 0.0001). **(F)** Images show motion analysis of Paxillin-GFP after TIRF imaging. Scale bar, 25 μm. **(G)** Western blots showing indicated protein levels in 690.cl2 cells after transfection with control (siCtr) and two individual Cdc42ep5 (siEp5) siRNAs. Right graphs show quantification of normalized p418-Src levels. Bars indicate mean ± SEM (*n*=5 independent experiments; one-way paired ANOVA, Dunnet’s test: *, *p* < 0.05; ***, *p* < 0.001).

To establish a causal connection between Cdc42ep5 filaments and actomyosin structures in 2D, Cdc42ep5 expression was silenced in 690.cl2 cells using two independent RNAi. Cdc42ep5 knockdown severely affected F-actin organization, as actin stress fibres were completely disrupted, especially at the perinuclear region (**Fig 3C**). In contrast, the meshwork pattern of fibrillary actin at the cell periphery was not affected, albeit actin in the lamellipodial areas appeared thinner and more condensed. This result showed that Cdc42ep5 was particularly required for stress fibre stabilization. Importantly, CDC42EP5-associated phenotypes were extensible to other cell types as similar results were obtained after depletion of CDC42EP5 in human melanoma cells WM266.4 and in murine embryonic fibroblasts (**Fig EV3C**). This data suggests that CDC42EP5 is a conserved regulator of the actomyosin cytoskeleton.

The actomyosin complex is critical for the growth and elongation of cell-ECM adhesions required for efficient cell migration, as it mediates the maturation of nascent focal complexes to FAs (27). This process requires myosin-mediated tension as well as the formation and maintenance of an F-actin network (28). The protein Paxillin is recruited early into focal complexes and plays a key scaffolding role at FAs. Its phosphorylation at Y31 and Y118 is crucial for FA formation (29). Our previous observations suggested that Cdc42ep5 may be part of the actomyosin machinery that drives the maturation of focal complexes to FAs in melanoma. To test this in more detail, we first investigated Cdc42ep5 association with FAs. Confocal microscopy analyses showed that Cdc42ep5 was present at actin fibres that connected with FAs but showed no clear localization at pY188-Paxillin areas (**Fig 3D**). Together, these data indicated that Cdc42ep5 is not part of the FA but localizes at high contractile F-actin filaments associated with FA, suggesting a potential role in FA maturation. To test this, FA formation and dynamics in 690.cl2 cells were studied after *Cdc42ep5* silencing. Using pY118-Paxillin staining, we observed that Cdc42ep5-depleted cells presented smaller and dot-like adhesions that localized closer to the cell periphery compared with control cells (**Fig 3E**), features that are characteristic of immature focal complexes (29). Zyxin is a FA protein that distinguishes mature FA from focal complexes as it is recruited later into the complex (29). Knockdown of Cdc42ep5 in 690.cl2 cells resulted in a complete loss of Zyxin-positive adhesions (**Fig EV3D**). Importantly, this defect was observed when cells were cultured both on glass and on fibronectin, ruling out any potential effect associated with defective cell adhesion. Furthermore, time-lapse imaging of Paxillin-GFP showed that Cdc42ep5 knockdown affected the elongation of FAs, which appeared static over time (**Movie EV4; Fig 3F**). As a result of deficient FA maturation, Cdc42ep5-depleted cells failed to activate FA downstream signalling (i.e. Src)(19) (**Fig 3G**) and presented defects in single cell migration in 2D (**Fig EV3E**). Interestingly, it was observed that Cdc42ep5 was the main Borg protein involved in the process. While silencing Cdc42ep1 and Cdc42ep3 marginally reduced pY418-Src levels, knocking-down Cdc42ep5 resulted in significant inhibition of Src activity (**Fig EV3F**). Together, these data indicated that in 2D melanoma cells Cdc42ep5 forms filamentous structures that overlap actin filaments, and promotes the formation of perinuclear actomyosin fibres leading to the maturation of FAs and increased cell motility.

### Sept9 promotes actomyosin activity, invasion and metastasis in melanoma

Borg proteins including Cdc42EP5 have been shown to primarily promote septin filament assembly (19, 30). In humans, septins are encoded by 13 different genes that encode multiple septin isoforms (10). Septins can associate with distinct actin networks, modulate their formation and facilitate myosin activation (11). In amoeboid T lymphocytes, it has been proposed that septins tune actomyosin forces during motility (14). To assess whether Cdc42EP5 was functioning via septins to regulate actomyosin function and invasion in melanoma we first analysed whether there was any particular septin associated with melanoma aggressiveness. Analyses of datasets of human material (31, 32) indicated that *SEPT9* was the only septin gene whose upregulation is associated with melanoma progression and dissemination (**Fig 4A; Fig EV4A and B**). These results pointed to a specific role of SEPT9 in melanoma invasion and metastasis that we sought to investigate further.

**Figure 4.**
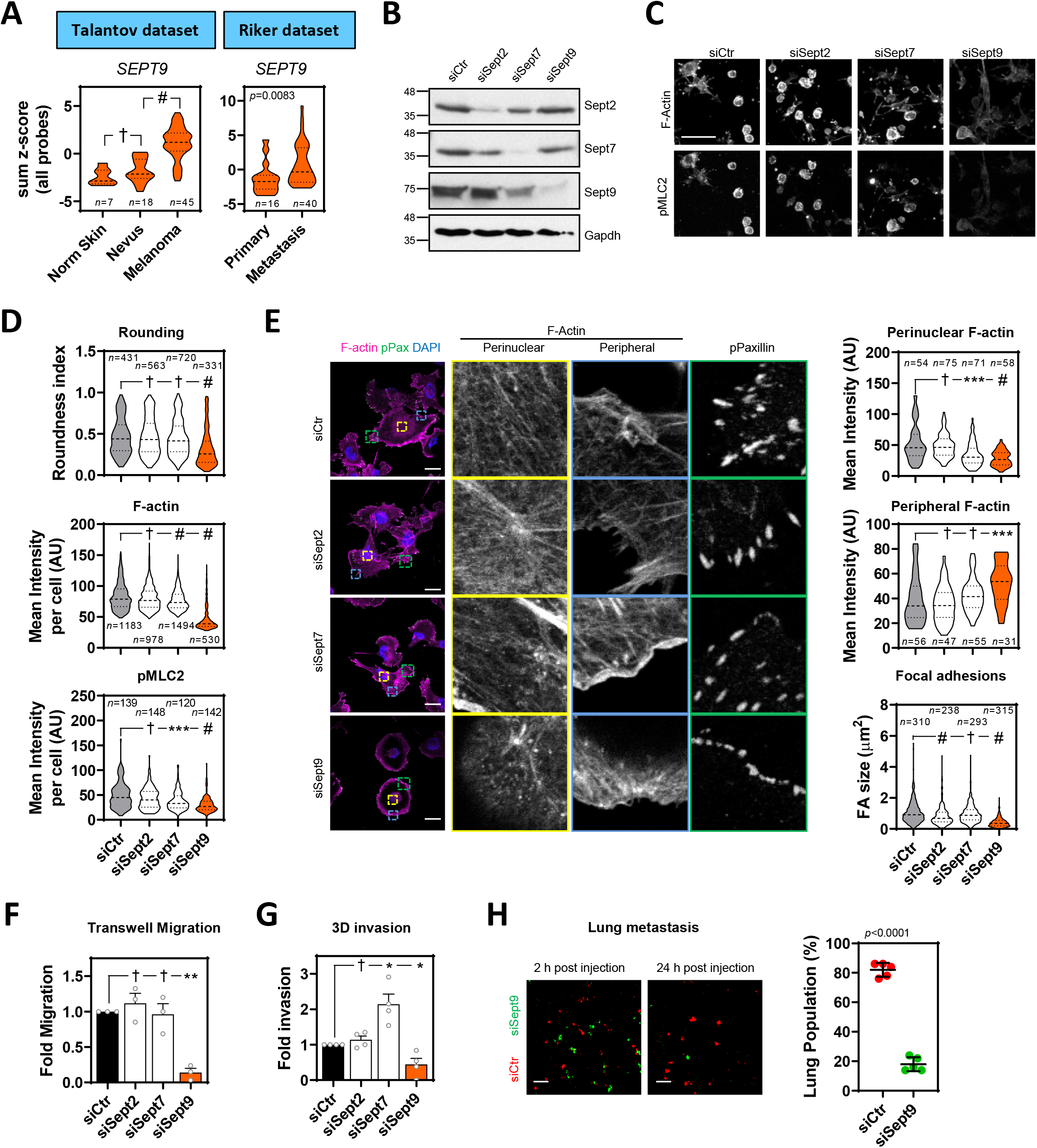
SEPT9 is required for actomyosin contractility and invasion in melanoma. **(A)** Violin plots show *SEPT9* expression in human tissues from normal skin, nevus and melanoma (*left*, Talantov dataset, GSE3189) and from primary melanoma and metastatic melanoma (*right*, Riker dataset, GSE7553); *n*, individual patients; Kruskal-Wallis and Dunn’s tests (Talantov) and Mann-Whitney test (Riker): †, not significant; #, *p* < 0.0001. **(B)** Western blots showing indicated protein levels in 690.cl2 cells after transfection with control (siCtr) and Sept2, Sept7 and Sept9 siRNAs. **(C)** Images show F-actin and pS19-MLC2 staining of parental 690.cl2 after transfection with control (siCtr), Sept2, Sept7 and Sept9 siRNAs and seeded on collagen-rich matrices. Scale bar, 75 μm. **(D)** Violin plots showing roundness index (*top*), F-actin mean intensity (*centre*) and pS19-MLC2 mean intensity (*bottom*) of individual cells from (C); *n*, individual cells; Kruskal-Wallis and Dunn’s tests: †, not significant; ***, *p* < 0.001; #, *p* < 0.0001. **(E)** Images show F-actin (magenta), pY118-Paxillin (green) and DAPI (cyan) staining in 690.cl2 cells after transfection with indicated siRNAs and seeded on glass. Right panels show indicated magnifications of F-actin staining in the perinuclear and peripheral regions, and pY118-Paxillin staining at the cell border. Scale bars, 25 μm. Violin plots show quantification of F-actin intensity in the perinuclear (*top*) and peripheral (*centre*) regions, and quantification of focal adhesion (FA) size (*bottom*); *n*, single regions from individual cells (*top* and *centre*) and individual FAs (*bottom*); Kruskal-Wallis and Dunn’s tests: †, not significant; ***, *p* < 0.001; #, *p* < 0.0001. **(F** and **G)** Graph shows fold migration ability (F) and fold invasion into collagen (G) of 690.cl2 cells after transfection with indicated siRNAs. Bars indicate mean ± SEM (*n*=3 [F] and 4 [G] independent experiments; paired *t* tests: †, not significant; *, *p* < 0.05; **, *p* < 0.01). **(H)** Images of mouse lungs at 2 and 24 h after tail injection with 690.cl2 cells transfected with control (siCtr-Red) or Sept9 (siSept9-Green) siRNA. Scale bar, 100 μm. Graph shows proportions (%) of siCtr and siSept9 cells within the lungs at 24 h. Lines represent mean ± SEM (*n*=5 mice; unpaired *t* test).

SEPT9 is the only member of the septin family that directly promotes actin polymerisation and crosslinking (12, 33). However, other septins such as SEPT2 and SEPT7 have been shown to co-localise with actin fibres and modulate their formation in non-malignant cells such as fibroblasts (19), lymphocytes (14) and endothelial cells (13), as well as breast cancer cells (34). To assess SEPT9 relevance in melanoma in comparison to other critical septins (i.e. SEPT2 and SEPT7), we proceeded to analyse the effect of perturbing Sept2, Sept7 and Sept9 function in 690.cl2 cells by specifically targeting their expression by RNAi. All individual treatments specifically decreased the expression of their targeted septin, albeit there was a slight reduction in Sept9 expression after depletion of Sept7 (**Fig 4B; Fig EV4C**). Only disruption of Sept9 expression affected both cell roundness and the generation of an actomyosin cortex (as read by pS19-MLC2 and F-actin levels) in 690.cl2 cells seeded on collagen-rich matrices (**Fig 4C and D; Fig EV4D**). Similar findings were observed after SEPT9 depletion in human melanoma WM266.4 cells (**Fig EV4E**). Analyses of cells in 2D culture showed that only depletion of Sept9 and, to a lesser extent Sept7, was affecting the generation of perinuclear F-actin fibres (**Fig 4E**). Similar to Cdc42ep5 depletion, silencing Sept9 led to a marked reduction in FA size(**Fig 4E**). Sept2-depleted cells also exhibited smaller FAs. In support of a specific role of Sept9 in controlling actomyosin contractility in melanoma, we observed that Sept9 silencing significantly reduced the abilities of parental 690.cl2 cells to migrate in transwell assays (**Fig 4F**). There was no significant change in the migration ability after Sept2 or Sept7 knockdown. In 3D collagen invasion assays, knockdown of Sept9 resulted in a significant decrease in invasion (**Fig 4G**). On the other hand, Sept2 depletion did not affect invasion whereas silencing Sept7 surprisingly increased the invasive potential of 690.cl2 cells. Importantly, these phenotypic and functional defects were associated with a reduced ability of 690.cl2 cells to metastasize in tail-vein assays after Sept9 depletion (**Fig 4H**). Altogether, this data points at SEPT9 as the key regulator of the actomyosin cytoskeleton during melanoma local invasion and distant metastatic colonisation.

### Cdc42ep5 promotes SEPT9 actin crosslinking leading to enhanced actomyosin activity

Having established a specific role for SEPT9 in regulating actomyosin contractility and invasion in melanoma, we next sought to investigate whether Cdc42EP5 was modulating SEPT9 function in these processes. 690cl2 cells in 2D cultures presented Cdc42ep5 co-localised with Sept9 filaments (**Fig 5A**). These filaments exhibited a clear overlap with F-actin stress fibres in the perinuclear region but not in the cell periphery. Similarly, Sept9 was also observed in the actin cortex with Cdc42ep5 and F-actin in amoeboid 690.cl2 cells seeded on collagen rich 3D matrices (**Fig 5B**). Importantly, modulating Cdc42ep5 expression amply affected the formation of Sept9 structures. Thus, we observed that silencing Cdc42ep5 reduced the formation of perinuclear Sept9 filaments and increased its cytosolic pool in 2D culture (**Fig 5C**). In 3D environments, Sept9 presented a sparse cytosolic pattern in 690.cl2^KO^-GFP, whereas stable reconstitution of Cdc42ep5 expression redistributed Sept9 to cortical regions cells (**Fig 5D**).

**Figure 5.**
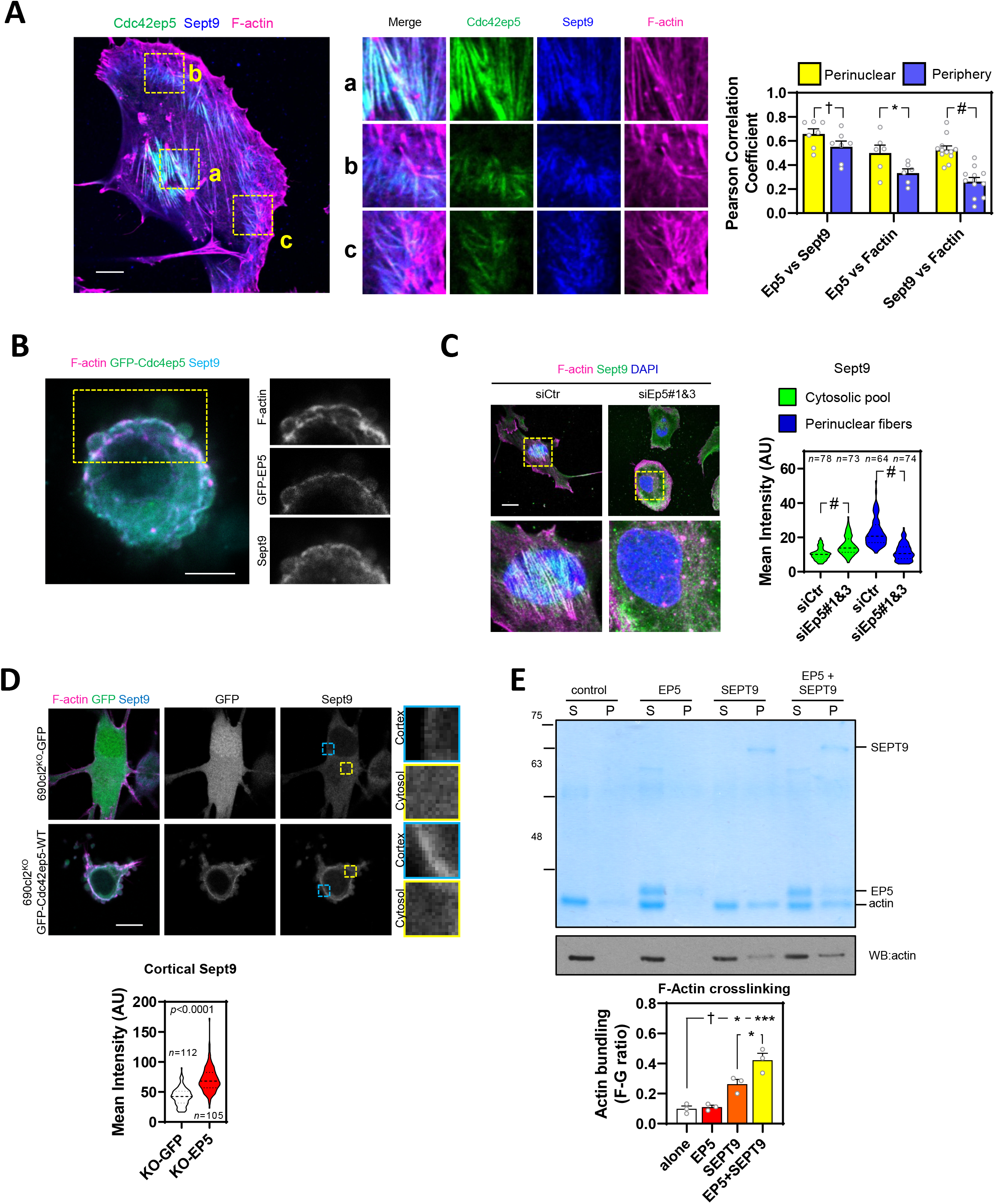
Cdc42ep5 associates with Sept9 and F-actin to promote F-actin crosslinking and the generation of actomyosin structures. **(A)** Left panel shows a 690.cl2 cell expressing gFP-Cdc42ep5 (green) and stained for Sept9 (blue) and F-actin (magenta). Scale bar, 10 μm. Middle panels show merged and individual channel magnifications of indicated areas: a, perinuclear; b and c, peripheral. Graph shows Person Correlation Coefficient of Cdc42ep5, Sept9 and F-actin in perinuclear and peripheral areas. Bars indicate mean ± SEM (*n*=7, 6 and 12 independent regions in individual cells; unpaired *t* tests: †, not significant; *, *p* < 0.05; #, *p* < 0.0001). **(B)** Image of 690.cl2 cell expressing GFP-Cdc42ep5 (green) seeded on collagen-rich matrices. Staining of F-actin (magenta) and Sept9 (cyan) and individual channel magnifications of indicated area are also shown. Scale bar, 10 μm. **(C)** Images show F-actin (magenta), Sept9 (green) and DAPI (blue) staining in 690.cl2 cells on glass after transfection with indicated siRNAs. Magnifications of perinuclear areas are shown. Scale bar, 20 μm. Violin plot shows quantifications of cytosolic and perinuclear intensity of Sept9 staining; *n*, individual cells; Mann-Whitney’s tests: †, not significant; ***, *p* < 0.001; #, *p* < 0.0001 **(D)** Images show F-actin (magenta), GFP (green) and Sept9 (cyan) of 690.cl2^KO^ cells expressing GFP or GFP-Cdc42ep5 and seeded on collagen-rich matrices. Scale bar, 10 μm. Single channels for GFP and Sept9 signals are shown. Panels on the right show magnifications of Sept9 signals in cortical and cytosolic areas. Violin plot at the bottom shows quantification of cortical Sept9 signal; *n*, independent regions in individual cells; Mann-Whitney’s test. **(E)** Top panel is a Coomassie-stained gel showing equal volumes of supernatant (S) and pellet (P) fractions from low-speed sedimentation of pre-polymerized actin filaments in the presence of recombinant Cdc42EP5 (1μM), SEPT9 (1μM) and both Cdc42eEP5 (1μM) and SEPT9 (1μM). Bottom panel shows an anti-Actin Western Blot of the same experiment. Graph shows the actin bundling coefficient (ratio of actin in pellet vs supernatant) in the indicated experimental points. Bars indicate mean ± SEM (*n*=3 experiments; one-way ANOVA, Tukey’s test: †, not significant; *, *p* < 0.05; ***, *p* < 0.001).

As opposed to other septins, SEPT9 is the only known septin that binds actin filaments directly and can promote the crosslinking of pre-polymerised actin filaments, even at suboptimal concentrations (12, 33)(**Fig EV5A**). Using low-speed actin sedimentation assays we confirmed that neither Cdc42EP5 nor SEPT6/7 induced F-actin crosslinking on their own (**Fig EV5B**). Interestingly, we observed that addition of recombinant Cdc42EP5 increased the amount of crosslinked F-actin induced by SEPT9 (**Fig 5E**). In addition, using F-actin polymerization assays and F-actin binding assays we confirmed previous findings describing the ability of recombinant SEPT9 to bind F-actin and promote actin polymerization (12, 33)(**Fig EV5C and D**); on the other hand, Cdc42EP5 exhibited no actin polymerization activities and a minimal interaction with F-actin.

### Sept9 is a crucial effector of Cdc42EP5 function in melanoma

These data suggested that Cdc42EP5 acted primarily by reinforcing the ability of SEPT9 to cross-link actin filaments into stress fibres and cortical actin, required for actomyosin contractility in 2D and 3D settings, respectively. To further validate this idea, we generated a Cdc42EP5 mutant defective in interaction with SEPT9 by mutating key residues in the BD3 domain of Cdc42ep5 (Cdc42ep5^GPS-AAA^ mutant, **Fig 6A, Fig EV6A**)(19, 30). We assessed the activity of this mutant in comparison to wild-type Cdc42ep5 (Cdc42ep5^WT^) by gain-of-function analyses in 690.cl2^KO^ (**Fig EV6B**). Contrary to Cdc42ep5^WT^ reconstitution, expression of Cdc42ep5^GPS-AAA^ in 690.cl2^KO^ cells in 2D did not induce the formation of perinuclear Sept9 networks (**Fig EV6C**), actin stress fibres or matured FAs (**Fig 6B**). In 3D, expression of Cdc42ep5^GPS-AAA^ in 690.cl2^KO^ did not affect cell rounding and pS19-MLC2 levels whereas Cdc42ep5^WT^ significantly increased these parameters (**Fig 6C; Fig EV6D and E**). Importantly, these phenotypes were concomitant with functional defects, as septin binding was absolutely required for Cdc42ep5-dependent transwell migration and collagen invasion (**Fig 6D and E**). Overall, these results confirm that Cdc42ep5 requires Sept9 interactions to promote actomyosin-dependent pro-invasive behaviours in melanoma.

**Figure 6.**
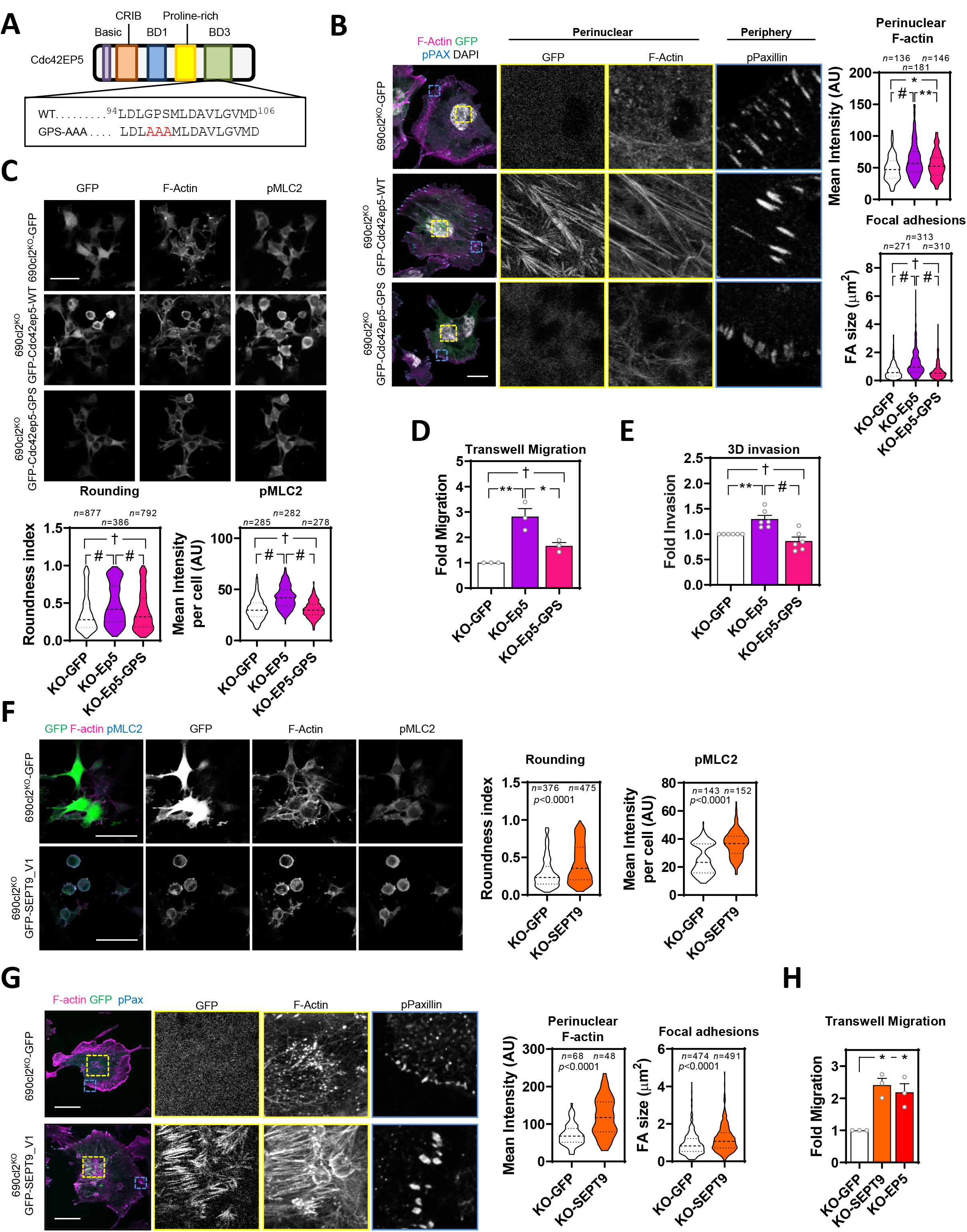
SEPT9 is a crucial effector of Cdc42EP5 function in melanoma. **(A)** Diagram showing the different domains in Cdc42ep5. Underneath, aminoacid sequence of BD3 segment in wild-type and septin-binding defective mutant (GPS-AAA). **(B)** Images show GFP (green), F-actin (magenta), pY118-Paxillin (cyan) and DAPI (grey) in 690.cl2^KO^ cells expressing GFP, GFP-Cdc42EP5^WT^ or GFP-Cdc42EP5^GPS-AAA^ and seeded on glass. Single channel magnifications of indicated areas are shown. Scale bar, 25 μm. Violin plots show quantification of perinuclear F-actin intensity (*top*) and focal adhesion size (*bottom*); *n*, independent regions in individual cells; one-way ANOVA Tukey’s test: †, not significant; *, *p* < 0.05; *, *p* < 0.05; #, *p* < 0.0001. **(C)** Images show GFP, F-actin and pS19-MLC2 in 690.cl2^KO^ cells expressing GFP, GFP-Cdc42EP5^WT^ or GFP-Cdc42EP5^GPS-AAA^ and seeded on collagen-rich matrices. Scale bar, 75 μm. Bottom graphs show violin plots for roundness index (*left*) and pS19-MLC2 mean intensity (*right*) of individual cells; *n*, individual cells; Kruskal-Wallis and Dunn’s tests: †, not significant; #, *p* < 0.0001. **(D and E)** Graphs show fold migration ability (D) and 3D collagen invasion (E) of 690.cl2^KO^ cells expressing GFP, GFP-Cdc42EP5^WT^ or GFP-Cdc42EP5^GPS-AAA^. Bars indicate mean ± SEM (*n*=3 [D] and 6 [E]; one-way ANOVA Tukey’s test: †, not significant; *, *p* < 0.05; **, *p* < 0.01; #, *p* < 0.0001). **(F)** Images show F-actin (magenta), GFP (green) and pS19-MLC2 (cyan) in 690.cl2^KO^ cells expressing GFP or GFP-SEPT9_V1 and seeded on collagen-rich matrices. Individual channels are also shown. Scale bar, 75 μm. Violin plots show roundness index (*left*) and pS19-MLC2 mean intensity (*right*) of individual cells; *n*, individual cells; Mann-Whitney test. **(G)** Images showing GFP (green), F-actin (magenta) and pY118-Paxilin (cyan) in 690.cl2^KO^ cells expressing GFP or GFP-SEPT9_V1 on glass. Right panels show indicated magnifications. Scale bar, 25 μm. Violin plots show quantification of F-actin intensity in the perinuclear region (*left*) and of focal adhesion (FA) size (*right*); *n*, single regions from individual cells (*left*) and individual FAs (*right*); Mann-Whitney test. **(H)** Graph shows fold migration ability of 690.cl2^KO^ cells expressing GFP, GFP-SEPT9_V1 or GFP-Cdc42ep5^WT^. Bars indicate mean ± SEM (*n*=3 experiments; paired *t* test: *, *p* < 0.05).

To further confirm that Sept9 was a crucial effector of Cdc42ep5 function in melanoma, we next investigated whether elevating Sept9 activity was sufficient to rescue the functional defects associated with loss of Cdc42ep5. For this, we used a SEPT9 isoform containing the N-terminal domain (SEPT9_V1), which has been shown to present enhanced activities (35). Thus, reconstituting Sept9 activity in 690.cl2^KO^ cells by ectopic expression of SEPT9_V1 was able to induce cell rounding and actomyosin activity in 3D culture (**Fig 6F; Fig EV 6F**). In addition, SEPT9_V1 expression reconstituted perinuclear actin fibre and FA formation in 690.cl2^KO^ cells (**Fig 6G**) and increased their migratory capabilities to similar levels as those obtained by Cdc42ep5 expression (**Fig 6H**). Overall these results illustrate a unique role for Sept9 in controlling actomyosin structure and function in melanoma, and place it as the key effector of Cdc42ep5 functions in regulating migration, invasion and metastasis.

## Discussion

Here we identify Cdc42EP5 as a new regulator of melanoma invasion and metastasis. We demonstrate that Cdc42EP5 forms filamentous structures that associate with F-actin and promotes the assembly of higher order actomyosin bundles in 2D and 3D. Contrary to other actin bundling regulators such as Fascin, alpha-Actinin and Filamin (36), Cdc42EP5 does not promote actin bundling directly but by controlling septin network reorganization. Thus, a septin-binding defective mutant is not able to affect actin structures or to confer migratory and invasive capabilities in melanoma.

Septins such as SEPT2, SEPT4, SEPT6, SEPT7 and SEPT9 have been shown to co-localise with actin most prominently in stress fibres, and their perturbation can alter actin organization (11). In particular, SEPT7 is proposed to be the essential septin in the formation of septin oligomers and filamentous structures (37). In addition, SEPT2 binds Myosin II directly and this interaction is proposed to promote actin:septin filament association and enhance MLC2 activation (38). In amoeboid lymphocytes, SEPT7 has been shown to modulate the actomyosin cytoskeleton in cortical contraction (14). SEPT9 occupies the terminal position in septin oligomers and has been proposed to be essential for filament formation, although perturbation of SEPT9 function only results in late abscission defects during cytokinesis but does not affect septin-dependent steps earlier in mitosis (39). On the contrary, SEPT2, 7, and 11 are required at the early stages of cytokinesis (40). Our orthogonal analyses of differential gene expression in clinical samples coupled to loss-of-function characterisation indicate that SEPT9 is the main septin effector of Cdc42EP5, which was further confirmed using *in vitro* analyses and epistatic approaches. In the context of melanoma, SEPT9 is essential for the generation of an actomyosin cortex in 3D and stress fibres in 2D. Altogether, it appears that SEPT9 functions in actomyosin networks independently of the formation of canonical SEPT2/6/7/9 oligomers. This specific role appears to be critical in the stabilisation of highly contractile actomyosin structures such as the cleavage furrow (40), stress fibres (12)(**Fig 4**) and actomyosin cortex (**Fig 4**), and may be related to the unique ability of SEPT9 to crosslink F-actin that is not shared by the rest of septins (12, 33). In agreement, we demonstrate mechanistically that Cdc42EP5 interacts with SEPT9 and potentiates its F-actin bundling activity required for the stabilisation of actomyosin networks

Actomyosin networks such as the actomyosin cortex in 3D and stress fibres in 2D are critical structures for exerting and resisting mechanical tension. In addition to the myosin motors required for generating force, these structures require additional levels of F-actin crosslinking to generate bundles capable of resisting the mechanical loads applied to them. Myosin II is the main actin-based contractile myosin motor that crosslinks actin filaments at the cell cortex and stress fibres and regulates cellular tension (9, 36). Accordingly, actomyosin activity in melanoma has primarily been shown to be exerted via Rho-ROCK modulation of myosin activity (2–5, 23), and is critical in modulating fast amoeboid invasion and metastasis (4, 23). We demonstrate that actomyosin function in melanoma is also dependent on the additional support provided by SEPT9 in those structures, which is modulated by the septin regulator Cdc42EP5. Thus, we observe both Cdc42EP5 and SEPT9 co-localising with actomyosin networks in 2D and 3D, and their silencing is associated with the destabilisation of F-actin structures and reduced actomyosin function. Thus, Cdc42EP5:SEPT9 filaments appear to complement Myosin II function and mechanically stabilise these structures. This new regulatory axis of actomyosin function in melanoma is required for invasion and metastasis.

Septins are increasingly associated with cytoskeletal regulation and cell migration, and potential roles in modulating cancer aggressiveness are emerging (11, 16). Determining their relevance in different cancers and understanding how they are regulated is therefore critical. We show a specific *SEPT9* upregulation in human melanoma and particularly in metastatic melanoma, which may underpin its functional relevance in this setting. In addition, we show that melanoma cells present characteristic SEPT9 networks that modulate their migratory behaviour by potentiating actomyosin activity. In addition, we identify a new regulatory mechanism of septin function in cancer. Thus, Cdc42ep5 is required for the assembly of SEPT9 structures at actomyosin bundles in 2D and 3D, possibly by enabling its correct positioning within the cell or within actin filaments. Thus, silencing of Cdc42ep5 in melanoma cells seeded in 2D leads to the disassembly of higher order Sept9 filaments, whereas in 3D results in reduced cortical Sept9 (**Fig 7**).

**Figure 7.**
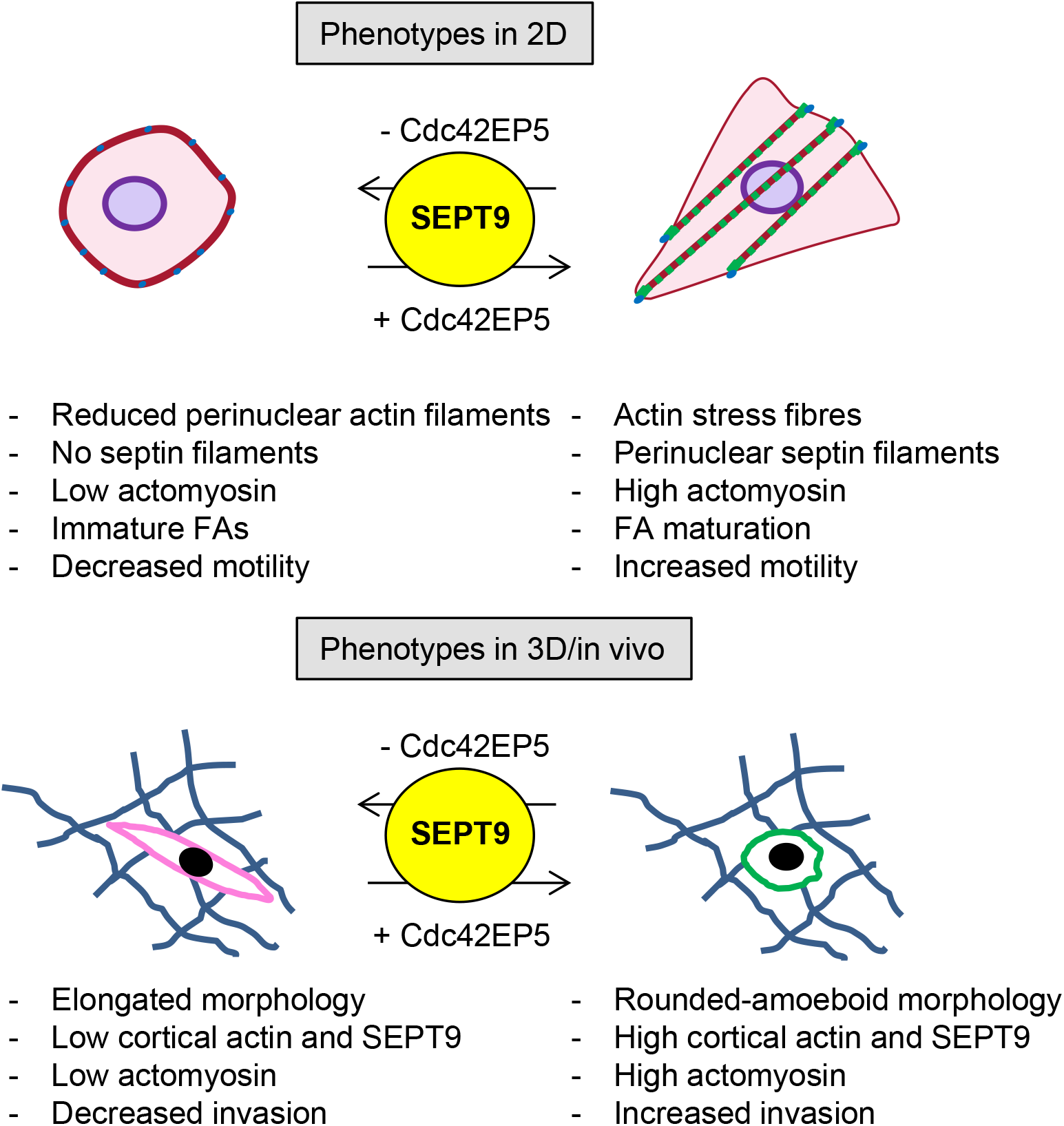
Cdc42EP5 is required for melanoma migration and invasion by promoting actomyosin contractility via SEPT9. Diagram summarizing the main findings of this study.

Melanoma cells can switch between mesenchymal and amoeboid modes of migration and this plasticity is a critical factor in the difficulties encountered in the clinic to target metastasis (41). Since actomyosin is required for both migratory modes, our results suggest that the Cdc42EP5-SEPT9 axis could potentially reduce both rounded-amoeboid and elongated-mesenchymal modes of invasion. Thus, cells lacking Cdc42EP5 or SEPT9 present defects in stress fibre formation and FA maturation which are generally required for mesenchymal migration. In agreement, silencing Cdc42ep5 resulted in a significant decrease in directed migration and motility in 2D, which favours elongated-mesenchymal behaviours. We also describe that CDC42EP5 is required for migration and invasion in a mesenchymal model of breast cancer such as MDA-MB-231. In breast cancer, SEPT9 has been reported to promote invasion by enhancing ECM degradation through metalloproteinase secretion (42), a hallmark of mesenchymal migratory behaviours. Similar results were obtained in a model of renal epithelial-mesenchymal transition, where SEPT9 was required for mesenchymal migration (12). Together, these findings suggest that the Cdc42EP5-SEPT9 axis may be an interesting node to target melanoma metastasis, as it might affect both modes of migration and circumvent the problem of melanoma cell plasticity (23).

Although the link between Borg proteins and cytoskeletal rearrangements is long standing (43), very little is known regarding their relevance in cancer and invasion. Here, we have performed for the first time a comparative study of the ability of different Borgs proteins to regulate cancer cell migration. Our analyses identify a unique role for CDC42EP5 in conferring pro-migratory and pro-invasive properties that is not consistently shared by other members of the family. This is surprising, as all Borgs share both regulatory and effector domains (18). We can speculate that specificity in this case is achieved via transcriptional regulation of these genes. Thus, we show an specific upregulation of *CDC42EP5* in rounded-amoeboid melanoma cells and describe a concomitant function of CDC42EP5 in modulating actomyosin activity in melanoma. This may explain its unique requirement for migration in confined environments such as collagen invasion assays or *in vivo* migration, as these processes rely significantly in actomyosin function. On the other hand, other Borgs may participate in modes of migration that require different cytoskeletal rearrangements. For example, CDC42EP4 function on normal mammary epithelia 2D migration was associated to enhanced filopodia formation (44). Still to be determined are the molecular features conferring this unique role to CDC42EP5. This may result from unique binding partners, structural characteristics or additional regulations that dictate its specific localization and/or function.

## Materials and Methods

### Cell lines

Murine melanoma 690.cl2 cells (a kind gift from Richard Marais, Cancer Research UK Manchester Institute, UK) were generated from spontaneous melanoma lesions from *Braf^V600E^* murine models (20, 21). WM266.4 human melanoma cells (a gift from Chris Bakal, The Institute of Cancer Research, UK) harbour a BrafV600D mutation and were isolated from a metastatic lesion (skin) of a female patient presented with malignant melanoma. MDA-MB-231 LM2 (a gift from Chris Bakal, The Institute of Cancer Research, UK) is a derivative of triple negative breast cancer cell model MDA-MB-231 that was selected for its ability to metastasize to lung tissue *in vivo*. Mouse embryonic fibroblasts (MEFs) were a kind gift from Afshan McCarthy (The Institute of Cancer Research, UK), and were kept at low passage and avoiding confluency. All these cell lines were cultured in DMEM (Sigma), GlutaMax (Gibco) and 10% foetal bovine serum (FBS) and incubated at 37°C in 5% CO2. A375P and A375M2 melanoma cells were from Richard Hynes (Massachusetts Institute of Technology, USA). CHL, SKMEL28, 501MEL, SKMEL2, SKMEL23, SBCL2, WM1361, WM1366 and WM3670 melanoma cells were from Richard Marais (Cancer Research UK Manchester Institute, UK). Cells were maintained in DMEM (RPMI for WM1361, SBCL2 and WM3670) containing 10% FBS and kept in culture for a maximum of 3–4 passages. All cell lines tested negative for mycoplasma infection with MycoAlert™ (Lonza).

### cDNA, RNAi and reagents

Murine pEGFP-Cdc42ep5 (N-terminal GFP) was a kind gift from Facundo Batista and Shweta Aggarwal (Francis Crick Institute, UK). This plasmid was used as a template to generate GFP-tagged mutant versions Cdc42ep5^GPS-AAA^ using In-Fusion cloning (Takara). To allow for lentiviral infection and stable expression, these cDNAs were all then subcloned into a pCSII-IRES-blasti vector backbone at NheI and BamHI restriction sites. The wild type version of Cdc42ep5 was also subcloned in-frame into pGEX backbone (pGEX-GST-Cdc42ep5) for generation of recombinant protein. Details of cloning strategy are available upon request. GFP-SEPT9_V1 plasmid was a kind gift from Cristina Montagna (Albert Einstein College of Medicine, USA). Paxillin-GFP was a kind gift from Chris Bakal (Institute of Cancer Research, UK). pCSII-IRES2-blasti-GFP and pCSII-IRES2-blasti-mCherry were a kind gift from Erik Sahai (Francis Crick Institute, UK). pnCS-Strep-SEPT6/7 is a bicistronic, spectinomycin resistant plasmid that expresses human SEPT6 and strep-SEPT7 in tandem; pET28-SEPT9 encodes a kanamycin resistant His-tagged version of human SEPT9 (45). Both plasmids were a kind gift from Elias Spiliotis (Drexel University, USA). siRNAs were purchased from Dharmacon and are listed in Table S1.

### Generation of CRISPR knock-out cell lines

The CRISPR plasmid U6gRNA-Cas9-2A-GFP containing a guide RNA targeting murine *Cdc42ep5* was purchased from Sigma (MM0000377239). Parental 690.cl2 cells were transfected with that plasmid using Lipofectamine (Life Technologies) following manufacturer’s instructions and GFP positive cells were single sorted into 96 well plates after 24 h. Individual cell clones were expanded and the *Cdc42ep5* locus targeted by CRISPR was sequenced and subjected to western blot analysis for knockout validation (690.cl2^KO^). *Cdc42ep5*-null clones were infected with GFP-Cdc42ep5-expressing lentivirus to generate 690.cl2^KO^ cells re-expressing wild-type Cdc42EP5 (690.cl2^KO^-GFP-Cdc42ep5) or the septin binding defective mutant of Cdc42EP5 (690.cl2^KO^-GFP-Cdc42ep5^GPS-AAA^). Alternatively 690.cl2^KO^ cells were infected with plain GFP expressing lentivirus to generate 690.cl2^KO^-GFP cells.

### Transfections

Cells were seeded at 75% confluency and transfected using RnaiMax (Life technologies) for siRNA (100 nM final concentration) and Lipofectamine 3000 (Life Technologies) for plasmids, following manufacturer’s instructions. Cell lines stably expressing cDNA (GFP-Cdc42ep5 or GFP/mCherry) were generated by lentiviral infection followed by blasticidin selection for 2 weeks (4 μg mL^−1^). Alternatively, fluorescent-labelled cells were sorted by FACS. For the generation of the non-virally transduced stable 690.cl2 cells expressing GFP, GFP-Paxillin or GFP-SEPT9_V1, parental cells were transfected with the relevant plasmids using Lipofectamine 3000, following manufacturer’s instructions. Next, 48 h after transfection, 400 μg mL^−1^ G418 (Sigma) was added as a selection agent to remove non-transfected cells. Cells were probed for the expression of plasmids based on GFP expression and kept at low passage.

### Proliferation assay

Cells were seeded in a 96 well in triplicate at 5,000 cells per well and grown in normal culture conditions. AlamarBlue assay was used according to manufacturer’s instructions to compare cell growth and viability at the indicated time points. Samples were run in quadruplicate and averaged. Values were normalized to values at day 0.

### Transwell Migration

5×10^4^ cells were suspended in serum-free media and seeded onto Transwell inserts with 8 μm pores (Corning) on a reservoir containing 10% FBS and 10 ng mL^−1^ TFGβ (Peprotech). After 48 h the cells on the lower chamber were fixed in 4% PFA for 15 min, stained with DAPI and imaged using a confocal microscope (Leica TCS SP8). A tile scan containing the entire well was acquired and number of cells (i.e. nuclei) quantified using Image J. Data is expressed as mean ± SEM (normalized to control cells).

### 3D inverted invasion assay

For 690.cl2: cells were suspended in 2.3 mg mL^−1^ rat tail collagen at 10^5^ cells mL^−1^. 100 μL aliquots were dispensed into 96-well flat bottom plates (Corning) pre-coated with 0.2% fatty acid free BSA (Sigma). Cells were spun to the bottom of the well by centrifugation at 300 × *g* for 5 min at 4°C, and incubated at 37°C in 5% CO2 for 1 h. Once collagen had polymerized, DMEM with 10% FBS and TGFβ (10 ng mL^−1^) was added on top of the collagen. After 24 h incubation at 37°C in 5% CO2, cells were fixed and stained for 4 h in 16% PFA solution containing 5 μg mL^−1^ Hoechst 33258 nuclear stain (Invitrogen). The plates were then imaged using an Operetta High Content Imaging System (PerkinElmer) at z-planes 0, 30, and 60 μm. Number of cells at each plane (i.e. nuclei) was quantified by the Harmony software package (PerkinElmer). The invasion index was calculated as the sum of the number of cells at 60 μm, divided by the total number of cells (at 0, 30 μm and 60 μm). The invasion index of each condition was then normalized to the respective control. Samples were run in triplicate and averaged. Data were expressed as mean ± SEM (normalized to control cells). Alternatively, cells were stained with phalloidin to generate 3D reconstructions of collagen invasion assays. Z-stacks (1 μm size) were acquired from bottom to top using a Leica SP8 confocal microscope and 3D reconstructions generated using LAS X software (Leica).

For WM266.4 and MDA-MB-231 cells: 100 μl of diluted 1:1 (Matrigel:PBS) mix was pipetted into Transwell inserts in a 24-well tissue culture plate and left to set for 30 min at 37°C. When the matrigel had set, the Transwell inserts were inverted and 5 × 10^4^ cells in normal media were pippeted onto the filter. The Transwell inserts were then carefully covered with the base of the 24-well tissue culture plate, making contact with each cell suspension droplet, and the plate incubated in the inverted state for 4 h to allow cell attachment. After this, plates were turned right-side-up and each Transwell insert washed with 3 × 1 ml serum-free DMEM and finally placed in a well containing serum-free DMEM. Full normal media (10% FBS) was gently pipetted on top of the set matrigel/PBS mixture and incubated for 5 d at 37°C in 5% CO2. After this, Transwell inserts were fixed in 4% PFA, stained with DAPI and phalloidin and imaged by confocal microscopy with a 20x objective with optical Z-sections scanned at 15-μm intervals moving up from the underside of the filter into the matrigel. The invasion index was calculated as the sum of the area of cells at 30 μm, divided by the total number of cells (at 0, 15 and 30 μm). Data were expressed as mean ± SEM (normalized to control cells).

### ECM-remodelling assay

To assess force-mediated matrix remodelling, 3×10^5^ cells were embedded in 120 μL of a 2.3 mg mL^−1^ rat tail collagen-1 gel in 24-well glass-bottom MatTek dishes and incubated at 37°C in 5% CO_2_ for 1 h. Once the gel was set, cells were maintained in normal culture conditions. Gel contraction was monitored daily by scanning the plates. The gel contraction value refers to the contraction observed after 2 days. To obtain the gel contraction value, the relative diameter of the well and the gel were measured using ImageJ software, and the percentage of contraction was calculated using the formula [100 × (well area – gel area) / well area]. Data were expressed as mean ± SEM.

### Generation of xenograft tumours and intravital imaging

All animals were kept in accordance with UK regulations under project license PPL80/2368. Six to eight week old CD1 nude mice were injected subcutaneously with 10^6^ 690.cl2^KO^-GFP or 690.cl2^KO^-Cdc42ep5 murine melanoma cells suspended in 100 μL of PBS:Matrigel (50:50). Tumour size was measured every other day using callipers. To calculate tumour volume the formula [V = (length × width^2^) / 2] was used. Intravital imaging using a Leica TCS SP8 microscope was performed when tumours reached 6-8 mm. Tumours were excited with an 880 nm pulsed Ti–Sapphire laser and emitted light acquired at 440 nm (collagen second harmonic generation, SHG) and 530 nm (GFP). During approximately 10-min intervals, 5 to 8 different regions were imaged simultaneously for 2 h for each tumour. In each region, a z-stack of 3 images (approximately 50 μm deep on average) was taken, resulting in a time lapse three-dimensional z series for analysis. Time lapse movies were processed and analysed using ImageJ, including a ‘3D drift correction’ script. This was achieved by converting the images obtained to hyperstacks and correcting for drift using the static SHG signal. The images generated were then processed to generate movies using the LAS X software. To generate coloured time projections, time-lapse movies were loaded into ImageJ and the function “Hyperstack>Temporal-Color Code” used. This generates motion analysis images by overlaying blue, green and red images from different time points. Distinct areas of colour indicate motile cells, whereas white areas indicate static regions. Once processed, images were assessed for cell movement, cell morphology and cell size. Moving cells were defined as those that moved 10 μm or more during the length of each movie, and the moving distance and resulting speed was determined using the LAS X software.

### Experimental metastasis assay

690.cl2 cells stably expressing GFP or mCherry were transfected with control or experimental siRNA (Cdc42ep5 or Sept9), respectively. 48 h post-transfection, Cherry and GFP cells were mixed in PBS at a ratio of 1:1, and 10^6^ cells (mixed population) were injected into the tail vein of CD1 nude mice. Mice were culled 2 and 24 h after injection, and lungs were dissected and placed in PBS. GFP and Cherry signal in fresh lungs were collected using a Leica SP8 confocal microscope (12 fields of view per lung). The area of GFP and Cherry cells was quantified using Volocity. For each mouse, the percentage of GFP and Cherry area of both lungs was averaged. Data were expressed as mean ± SEM from at least 4 independent animals.

### RNA isolation and qRT-PCR

RNA was isolated using RNeasy Kit (Qiagen). Reverse transcription was performed using Precision NanoScript 2 Reverse-Transcription-kit (PrimerDesign) and qPCR using PrecisionPLUS 2x qPCR MasterMix with ROX and SybrGreen (PrimerDesign). Expression levels of indicated genes were normalized to the expression of Gapdh and Rplp1. Sequences of the oligonucleotides used for qRT-PCR are described in Table S2.

### Co-Immunoprecipitation

690.cl2 cells expressing the GFP-tagged plasmids of interest were grown on 150 mm petri dishes and lysed in lysis buffer: 50mM Tris-HCl pH 7.5, 150mM NaCl, 1% (v/v) Triton-X-100, 10% (v/v) glycerol, 2mM EDTA, 25mM NaF and 2mM NaH_2_PO_4_. The resultant lysates were first pre-cleared using IgG-conjugated Protein G beads (ASD), then incubated with the specific antibodies for 2 h at 4°C and then incubated with Protein G beads for 1 h at 4°C with gentle mixing. Beads were then washed 4 times with lysis buffer and eluted with 20 μL of 2X SDS sample buffer.

### Generation of recombinant proteins

*Escherichia coli* BL21(DE3) bacteria transformed with pGEX-GST-Cdc42ep5, pnCS-Strep-SEPT6/7 or pET-28a-His-SEPT9 were grown in LB medium at 37°C with ampicillin, spectinomycin and kanamycin, respectively. Protein expression was induced at an OD600 of 0.8 with 0.5 mM IPTG for 16 hr at 18°C, harvested by centrifugation and suspended in 50 mM Tris-HCl pH 8, 300 mM NaCl, 1 mM PMSF and 5 mM benzamidine. Bacteria were sonicated 4 times on ice (amplitude 20%, 15 s on, 45 s off) and centrifuged at 20,000 × *g* for 30 min at 4°C. Supernatants were collected and incubated with 400 μL of the respective beads (Glutathione sepharose, Strep-Tactin or NiNTA agarose) for 1 hr. Beads were washed 5 times with 2CVs lysis buffer and eluted with 2 CVs of 50 mM Tris-HCl pH 8, 300 mM KCl and 5 mM MgCl_2_, supplemented with imidazole (250 mM), glutathione (10 mM) or desthiobiotin (2.5 mM). The eluted proteins were further purified by size exclusion chromatography on a Superdex 200 Increase 10/300 GL column (GE Healthcare) in 50 mM Tris-HCl pH 8, 300 mM KCl, 5 mM MgCl_2_ and 5 mM DTT. The concentration of all purified proteins was determined by the Bradford assay.

### In vitro Actin assays

Human non muscle actin (Cytoskeleton, Inc.) in general actin buffer (5 mM Tris-HCl [pH 8.0] and 0.2 mM CaCl2, supplemented with 0.2 mM ATP) was polymerized using actin polymerization buffer (50 mM KCl, 2 mM MgCl_2_, and 1 mM ATP) at 24°C for 1 hr to generate F-actin. This stock of F-actin was used for the bundling and binding assay. For the bundling assay, indicated concentrations of the different recombinant proteins (GST-Cdc42ep5, His-SEPT9 or Strep-SEPT6/7) were incubated with 2 μM F-actin for 1 hr at 24°C. Samples were centrifuged at 14,000 × *g* for 1 hr at 24°C. For the F-actin binding assay, the reaction of actin and the different recombinant proteins was centrifuged at 150,000 × *g* for 1.5 hr at 24°C. For the actin polymerization assay, actin was incubated on ice for 1 hr with general actin buffer to generate the G-actin stock. This stock was then mixed with the actin polymerization buffer and the indicated concentrations of the recombinant proteins at 24°C for 30 min. Samples were centrifuged at 150,000 × *g* for 1.5 hours at 24°C. In all cases, equal volumes of the supernatant and pellet fractions were resolved using a 12% SDS-PAGE gel, stained with Coomassie Brilliant Blue and processed by Western blotting (for GST, His and Actin) if indicated.

### Western Blotting

Protein lysates and immunoprecipitants were processed following standard procedures. Enhanced chemi-luminescence signal was acquired using an Azure Biosystems c600. Exposures within the dynamic range were quantified by densitometry using ImageJ. Antibody description and working dilutions can be found in Table S3.

### Immunofluorescence

Cells were seeded on glass bottom 24 well plates (MatTek). Where indicated, cells were seeded in glass bottom plates covered with 10 μg mL^−1^ Fibronectin. For analysis of 3D morphology, cells were seeded on top of 2.3 mg mL^−1^ collagen-I gel over a glass bottom dish (MatTek) in medium and allowed to adhere for 24 h. Cells were fixed in 4% PFA and permeabilised in PBS with 0.2% Triton-X. Where indicated (i.e. septin staining) cells were fixed in ice-cold methanol for 15 min and permeabilised in PBS with 0.2% Triton-X. Samples were blocked in 3% BSA with 0.1% PBS Tween (PBST) for 3 h. The primary antibodies (Table S3) were diluted in 3% BSA in PBST for 2 h. The wells were then washed 3×10 minutes in 3% BSA PBST, followed by the addition of the appropriate secondary (Alexa Fluor, Invitrogen). After 3 washes of 15 min in PBS, samples were mounted and analysed using a Leica SP8 confocal microscope. For analyses of fluorescence intensity differences, microscope settings were kept constant and independent replicates imaged on the same day.

### Time-lapse analysis and cell migration in 2D

Cells were seeded at low density and imaged for 18 h using bright field time lapse microscopy at 20x magnification. Individual cells were tracked using MTrackJ Image J plugin.

### Microscopy and Image analysis

For morphological analyses of cells grown over collagen, a minimum of 5 images were taken of cells per experiment using a 20x water objective on an inverted confocal microscope (Leica TCS SP8). Using phalloidin staining and Image J software, individual cells were selected and the roundness coefficient obtained using the Circularity function (4pi[area/perimeter^2]). For analysis of single cell F-Actin/pS19-MLC2/GFP intensity, individual cells were selected using Image J software and the mean fluorescence intensity of each channel per cell. Volocity was used for analysis of SEPT9 cytosolic/cortical intensity; areas in the centre of the cell (excluding the nucleus) were determined for the cytosolic region, and small regions towards the cell periphery (based on F-actin staining) were chosen to determine the cortical region of the cell. Cytosolic/cortical areas between samples were kept constant. The mean fluorescence intensity of Sept9 within individual areas was determined by the software.

For analysis of actin/septin/Cdc42ep5 structures, cells were stained for F-Actin/Sept9 and imaged at the basal plane using a 63x oil immersion lens. The basal perinuclear region of individual cells (based on the DAPI signal) was identified in each cell as the perinuclear region; the peripheral area was determined as a region close to the cell border and not in the vicinity of the nucleus. The fluorescence intensity of individual channels was determined using Volocity in the different areas.

For analysis of FA size, cells were stained with pY118-Paxillin antibody, imaged using a 63x oil immersion lens and analysed using Volocity to determine individual FA area. To determine the maturity of focal adhesions, Zyxin staining was used. A minimum of 5 fields of view per repetition was taken using a 63x oil objective, and cells were manually scored based on the presence or absence of Zyxin adhesions.

To determine the dynamics of Cdc42ep5 and F-actin, parental 690.cl2 cells were co-transfected with GFP-Cdc42ep5^WT^ and MARS-LifeAct. Cells were imaged using total internal reflectance microscopy (TIRF, 3i Imaging Solutions) for 60 s, using a 40x objective. To determine FA dynamics, a stable 690.cl2-GFP-Paxillin cell line was used. Cells were transfected with either control or RNAi targeting Cdc42ep5. TIRF microscopy was used to visualize adhesion dynamics. Cells were imaged for 10 min (1 frame every 30 s) using a 40x objective. To generate coloured time projections, time-lapse movies were loaded into ImageJ and the function “Hyperstack>Temporal-Color Code” used.

For co-localization analyses, cells were stained and imaged as before and processed using Image J software using the “Coloc2 plugin” (Pearson’s R value no threshold). For each individual cell, three independent ROI (perinuclear, peripheral or encompassing a focal adhesion) were analysed and the average value used as the Person’s Coefficient in that cell. Image acquisition and post-processing settings, and ROI size were maintained constant for each condition.

### Analysis of clinical datasets of melanoma

Gene expression data from Talantov (GSE3189)(31) and Riker (GSE7553)(32) studies were retrieved from NCBI GEO. Gene expression values for septin genes were calculated as a combination of all probes available for each gene. Briefly, all probes capturing septin gene expression on individual samples were z-score normalized. Final values of expression for individual septin genes in each sample were calculated by summing the z-scores of all probes available for each gene. *CDC42EP5* and some septin genes (i.e. *SEPT1*, *SEPT3*, *SEPT12* and *SEPT14* for Talantov, and *SEPT14* for Riker) were not probed in these analyses and therefore were not included in the analyses.

### Statistical analysis

Statistical analyses were performed using GraphPad Prism (GraphPad Software, Inc.). When *n* permitted, values were tested for Gaussian distribution using the D’Agostino-Pearson normality test. For Gaussian distributions, paired or unpaired two-tailed Student’s t-test and one-way ANOVA with Tukey post-test (for multiple comparisons) were performed. Following the software recommendations, Geisser-Grenhouse correction and Dennett’s multiple comparisons test were applied in paired one-way ANOVA. For non-Gaussian distributions, Mann-Whitney’s test and Kruskal-Wallis test with Dunn’s post-test (for multiple comparisons) were performed. Unless stated otherwise, mean values and standard errors (SEM) are shown. In violin plots were generated using GraphPad; in addition to showing the distribution, thick dashed lines represent the median whereas thin dashed lines represent the upper and lower quartiles. *P* values of less than 0.05 were considered statistically significant: *, *P* < 0.05; **, *P* < 0.01; ***, *P* < 0.001; #, *P* < 0.0001; †, non-significant. In each graph, the number of individual experimental points is described.

## Acknowledgments

**General**: We thank Facundo Bastista, Elias Spiliotis, Erik Sahai, Cristina Montagna and Chris Bakal for providing us with plasmids; Chris Bakal, Afshan McCarthy, Richard Hynes, Amine Sadok and Richard Marais for providing us with cell lines; Fredrik Wallberg (Light Microscopy Unit at The Institute of Cancer Research) and Victor Campa (Microscopy Unit at Instituto de Biomedicina y Biotecnología de Cantabria) for assistance; members of the Biological Services Unit at The Institute of Cancer Research for help with mouse experiments. We also thank present and past lab members and Chris Bakal for help and advice throughout this work.

## Author contributions

FC conceived the study. AJF and FC designed, performed and analysed the experiments, with help from JLO and VSM. JR generated recombinant proteins and performed *in vitro* analyses with help from ML. FC wrote the manuscript with essential contribution by AJF and VSM.

## Data and materials availability

The datasets and materials generated and/or analysed during the current study are available from the corresponding author upon reasonable request. Materials may be subjected to MTA.

**Figure EV1. Cdc42EP5 is required for melanoma migration and invasion.**

**(A)** Efficacy of RNAi silencing of Borg genes in 690.cl2 cells. Graph shows fold normalized expression of Cdc42ep1-5 (Ep1-5) against control cells (siCtr) cells when individual genes were targeted (siEp1-5). Bars indicate mean (*n*=2 independent experiments).

**(B)** Graph shows quantification of Cdc42ep5 protein levels in 690.cl2 cells after transfection with control (siCtr) or two individual Cdc42ep5 (siEp5) siRNA. Bars indicate mean ± SEM (*n*=3 experiments; one-way paired ANOVA Dunnet’s test; *, *p* < 0.05; **, *p* < 0.01). Representative Western Blot is shown in Fig 1C.

**(C)** Diagram showing the targeting sequence for endogenous Cdc42ep5 CRISPR/CAS9 knock-out in murine wild-type 690.cl2 cells. Underneath, sequence of the same locus in the 690.cl2^KO^ cells.

**(D)** Western blot showing Cdc42ep5, GFP and Gapdh expression in parental 690.cl2 cells (wild-type) and in 690.cl2^KO^ cells expressing GFP or GFP-Cdc42ep5.

**(E)** Graph shows fold migration ability of wild-type 690.cl2 (WT) cells compared to 690.cl2^KO^ cells expressing GFP or GFP-Cdc42ep5. Bars indicate mean ± SEM (*n*=3 experiments; one-way paired ANOVA Tukey’s test; *, *p* < 0.05; **, *p* < 0.01).

**(F)** Graph shows migration ability of parental 690.cl2 cells ectopically expressing GFP or GFP-Cdc42EP5. Bars indicate mean ± SEM (*n*=3 experiments; paired *t* test).

**(G)** Graph shows fold normalized expression of Cdc42EP5 in human WM266.4 and MDA-MB-231 cells after transfection with control (siCtr) and Cdc42EP5 (siEP5) siRNAs. Bars indicate mean (*n*=2 independent experiments).

**(H and I)** Graphs show fold migration ability (H) and 3D invasion (I) of WM266.4 and MDA-MB-231 cells after transfection with control (siCtr) and Cdc42EP5 (siEP5) siRNAs. Bars indicate mean ± SEM (*n*=3 experiments [H] and 4 [I]; paired *t* tests).

**(J)** Cell proliferation curves of 690.cl2^KO^ cells ectopically expressing GFP (KO) or GFP-Cdc42EP5 (KO-EP5). Lines indicate mean ± SEM (n=3 experiments).

**(K)** Graph shows quantification of volumes at day 7 post-injection of subcutaneous tumours induced by injection of 690.cl2^KO^ cells expressing GFP or GFP-Cdc42ep5. Bars indicate mean ± SEM (*n*=6 mice; paired *t* test).

**Figure EV2. Cdc42EP5 promotes actomyosin function in collagen-rich matrices.**

**(A)** Images of 690.cl2 cells expressing GFP-Cdc42ep5 (green) and seeded on collagen-rich matrices. Merged and single greyscale channels also show F-actin (magenta) and pS19-MLC2 (cyan) staining. Scale bar, 50 μm. Violin plots show pS19-MLC2 (*left*) and F-actin (*right*) mean intensity from individual cells in 690.cl2 cells expressing different levels of Cdc42ep5 (high or low); *n*, individual cells; Mann-Whitney test.

**(B)** Top panels represent the different shape of individual 690.cl2 cells and their respective roundness index. Left graph shows a violin plot of the roundness index of individual cells in 690.cl2 cells expressing different levels of Cdc42ep5 (high or low); *n*, individual cells; Mann-Whitney test. Right graph shows the relative frequencies of roundness indexes in Cdc42ep5^hi^ and Cdc42ep5^lo^ 690.cl2 cells.

**(C)** Higher magnification of a rounded-amoeboid 690.cl2 cell on a collagen-rich matrix expressing GFP-Cdc42ep5 (green) and stained for F-actin (magenta). Zoom up area of cell cortex and blebbing region is shown. Scale bar, 5 μm.

**(D)** Graph shows the relative frequencies of roundness indexes in 690.cl2 cells after transfection with control (siCtr) or two individual Cdc42ep5 (siEp5) siRNA. Additional representation of Figure 2C.

**(E)** Graph shows the percentage of cells within 3 different ranges of roundness, from elongated (<0.4) to rounded (>0.6) in in 690.cl2 cells after transfection with control (siCtr) or two individual Cdc42ep5 (siEp5) siRNA. Bars represent mean. Additional representation of Figure 2C and EV2D.

**(F)** Graphs show expression of *Cdc42EP1, Cdc42EP2, Cdc42EP3, Cdc42EP4* normalized to *GAPDH* expression in human melanoma cell lines with increasing rounding coefficients. Person correlation coefficient (r), statistical significance (P) and linear regression (red line) are shown. Each point in the graph represents the mean value of 3 independent experiments.

**(G)** Western blot of indicated proteins in 690.cl2 cells after transfection with control (siCtr) and siRNAs targeting individual Borg genes (siEp1-5). Graph shows the quantification of normalized pS19-MLC2 levels in the different experimental points. Bars indicate mean ± SEM (*n*=7 experiments; one-way paired ANOVA, Dunnet’s test: †, not significant; *, *p* < 0.05).

**Figure EV3. Cdc42EP5 modulates cytoskeletal features.**

**(A)** Image showing acetylated tubulin (red), GFP (green) and DAPI (blue) in 690.cl2 expressing GFP-Cdc2ep5 on glass. Right panels show single greyscale channel magnifications of the indicated area. Scale bar, 25 μm.

**(B)** Images show F-actin (magenta), GFP (green), pS19-MLC2 (cyan) and DAPI (grey) signal of 690.cl2 cells stably expressing GFP-Cdc42ep5 and seeded on glass. Top panels show merged channel images of a protrusion (*left*) and perinuclear areas (basal, *middle panel*; apical, *right panel*). Bottom right inserts show whole-cell images. Lower panels show individual greyscale channels for the indicated regions and signals. Scale bars, 10 μm.

**(C)** Images show F-actin (magenta) and DAPI (blue) staining of mouse embryonic fibroblasts (MEFs, *top panels*) and human melanoma cells (WM266.4, *lower panels*) seeded on glass after transfection with control (siCtr) and Cdc42EP5 (siEP5) siRNA. Greyscale magnifications of F-actin staining in indicated perinuclear areas are shown. Scale bar, 25 μm. Violin plot shows perinuclear F-actin intensity in the indicated experimental points and cell lines; *n*, single regions from individual cells; Mann-Whitney test: #, *p* < 0.0001.

**(D)** Images of 690.cl2 cells after transfection with control (siCtr) and Cdc42ep5 (siEp5#1&3) siRNA and seeded on glass. Images show F-actin (magenta), Zyxin (green) and DAPI (blue) staining. Right panels show greyscale Zyxin channel magnifications. Scale bar, 25 μm. Graph shows percentage of cells presenting Zyxin positive focal adhesions for cells seeded on glass or on fibronectin. Bars indicate mean ± SEM (*n*=4 experiments; unpaired *t* tests: #, *p* < 0.0001).

**(E)** Plots show single cell trajectories of 690.cl2 cells after transfection with control (siCtr) and two individual Cdc42EP5 (siEP5#1 and siEP5#3) siRNAs and seeded on 2D surfaces. Violin plot shows speed of individual cells (*n*, individual cells; Kruskal-Wallis and Dunn’s tests: *, *p* < 0.05; #, *p* < 0.0001).

**(F)** Western blots show pY438-Src and Tubulin levels in 690.cl2 cells after transfection with control (siCtr) and siRNAs targeting individual Borg genes (siEp1-5). Graph shows the quantification of normalized pY418-Src levels in the different experimental points. Bars indicate mean ± SEM (*n*=5 experiments; one-way paired ANOVA, Dunnet’s test: †, not significant; ***, *p* < 0.001).

**Figure EV4. SEPT9 is associated with melanoma aggressiveness.**

**(A and B)** Violin plots showing the expression of available septin genes in human tissues from normal skin, nevus and melanoma from Talantov dataset-GSE3189 (A), and from primary melanoma and metastatic melanoma from Riker dataset - GSE7553 (B); *n*, individual patients; Kruskal-Wallis and Dunn’s tests (A) and Mann-Whitney tests (B): †, not significant; *, *p* < 0.05; **, *p* < 0.01; ***, *p* < 0.001; #, *p* < 0.0001.

**(C)** Graph showing quantification of normalized protein expression levels of Sept2, Sept7 and Sept9 in 690.cl2 cells after transfection with control (siCtr) or Sept2, Sept7 or Sept9 siRNAs. Bars indicate mean± SEM (*n*=3 experiments; two-way ANOVA Dunnet’s test: †, not significant; *, *p* < 0.05). Representative Western Blot is shown in Fig 4B.

**(D)** Graph shows the relative frequencies of roundness indexes in 690.cl2 cells after transfection with control (siCtr) or Sept2, Sept7 or Sept9 siRNAs and seeded on collagen-rich matrices. Additional representation of Figure 4D.

**(E)** Top panels show morphologies of WM266.4 after transfection with control (siCtr) or SEPT9 (siSEPT9) siRNA and seeded on top of collagen-rich matrices. Scale bar, 20 μm. Left bottom graph shows roundness index quantification of individual cells (*n*, indicates individual cells; Mann-Whitney test). Right bottom graph shows the relative frequencies of roundness indexes.

**Figure EV5. Cdc42EP5 potentiates the F-actin bundling activities of SEPT9.**

**(A)** Coomassie-stained gel showing equal volumes of supernatant (S) and pellet (P) fractions from low-speed sedimentation of pre-polymerized actin filaments in the presence of different concentrations of recombinant SEPT9 (as indicated).

**(B)** Coomassie-stained gel showing equal volumes of supernatant (S) and pellet (P) fractions from low-speed sedimentation of pre-polymerized actin filaments in the presence of recombinant Cdc42EP5 (1μM), SEPT6/7 (1μM) and both Cdc42EP5 (1μM) and SEPT6/7 (1μM).

**(C)** Top panel is a Coomassie-stained gel showing equal volumes of supernatant (S) and pellet (P) fractions from high-speed sedimentation assays (F-actin binding assays) in the presence of recombinant Cdc42EP5 (1μM) and SEPT9 (1μM). Bottom panel shows anti-Actin, anti GST (Cdc42EP5) and anti-His (SEPT9) Western blots of the same experiment. SEPT9 only appears in the pellet (high interaction with F-actin) whereas Cdc42EP5 is primarily found in the supernatant (low interaction with F-actin).

**(D)** Top panel is a Coomassie-stained gel showing equal volumes of supernatant (S) and pellet (P) fractions from an actin polymerization assay in the presence of recombinant Cdc42EP5 (1M), SEPT9 (1M) and both Cdc42EP5 (1μM) and SEPT9 (1μM). Bottom panel shows an anti-Actin Western blot of the same experiment. Graph shows the actin polimerisation coefficient (ratio of actin in pellet vs supernatant) in the indicated experimental points. Bars indicate mean ± SEM (*n*=3 experiments; one-way ANOVA, Tukey’s test: †, not significant; *, *p* < 0.05).

**Figure EV6. Cdc42EP5 interacts with SEPT9 to promote actomyosin activity.**

**(A)** Western blot showing Sept9 and GFP levels in total lysates (input) and anti-GFP immunoprecipitates (IP:GFP) in 690.cl2 cells ectopically expressing GFP, GFP-Cdc42ep5^WT^ and GFP-Cdc42ep5^GPS-AAA^.

**(B)** Western blot showing Cdc42ep5 and Gapdh expression in parental 690.cl2 cells (wild-type) and in 690.cl2^KO^ cells expressing GFP, GFP-Cdc42ep5^WT^ or GFP-Cdc42ep5^GPS-AAA^.

**(C)** Images show GFP (green), Sept9 (blue) and F-actin (magenta) in 690.cl2^KO^ cells expressing GFP, GFP-Cdc42EP5^WT^ or GFP-Cdc42EP5^GPS-AAA^ and seeded on glass. Bottom panels are merged and single channel magnifications of perinuclear areas. Scale bar, 20 μm. Violin plot in the right shows perinuclear Sept9 mean intensity in the indicated cells (*n*, individual cells; Kruskal-Wallis and Dunn’s tests: †, not significant; #, *p* < 0.0001).

**(D)** Graph shows the relative frequencies of roundness indexes in 690.cl2^KO^ cells expressing GFP, GFP-Cdc42EP5^WT^ or GFP-Cdc42EP5^GPS-AAA^ and seeded on collagen-rich matrices. Additional representation of Figure 6C.

**(E)** Western blot showing pS19-MLC2 and Sept2 expression in 690.cl2^KO^ cells expressing GFP, GFP-Cdc42ep5^WT^ or GFP-Cdc42ep5^GPS-AAA^. Graph shows the normalized pS19-MLC2 levels. Bars indicate mean ± SEM (*n*=3 experiments; one-way ANOVA, Tukey’s test: †, not significant; *, *p* < 0.05; **, *p* < 0.01).

**(F)** Graph shows the relative frequencies of roundness indexes in 690.cl2^KO^ cells expressing GFP or GFP-SEPT9_V1 and seeded on collagen-rich matrices. Additional representation of Figure 6F.

## SUPPLEMENTARY MATERIAL

**Supplementary Table 1. siRNAs:** Name (target), species, sequence and catalogue number of the single siRNAs used in the study.

**Supplementary Table 2. Primers**. Names and sequences (forward and reverse) of the paired oligos used in this study for qRT-PCR. The name contains the target gene; h stands for human, m for murine; F for forward and R for Reverse.

**Supplementary Table 3. Antibodies**. Name, company, catalogue number and working dilutions of all the antibodies used in the study.

**Movie EV1. Intravital imaging of 690.cl2 cells expressing Cdc42EP5 in a living tumour.** 690.cl2-KO cells ectopically expressing GFP-Cdc42EP5 (green) were grown subcutaneously and imaged in vivo using two photon microscopy, which enable the acquisition of second harmonic signal (collagen fibres, magenta). Images were acquired every 14 min for a total of 2h 45min. Scale bar, Scale bar, 100 μm. Middle panels show magnifications of indicated areas showing fast amoeboid movement into the matrix. Right panels represent motion analysis images of the same area generated by overlaying blue, green and red images from different time points. Distinct areas of colour indicate motile cells appear, whereas static regions appear white.

**Movie EV2. Intravital imaging of 690.cl2 lacking Cdc42EP5 in a living tumour.** 690.cl2-KO cells ectopically expressing GFP (green) were grown subcutaneously and imaged in vivo using two photon microscopy, which enable the acquisition of second harmonic signal (collagen fibres, magenta). Images were acquired every 14 min for a total of 2h 45min. Scale bar, 100 μm. Middle panels show magnifications of indicated areas showing slow protrusive movement of cells and elongated morphology. Right panels represent motion analysis images of the same area generated by overlaying blue, green and red images from different time points. Distinct areas of colour indicate motile cells appear, whereas static regions appear white

**Movie EV3. Dynamic association between F-actin and Cdc42EP5.** Time-lapse movie of 690.cl2 cell expressing LIFEACT-RFP (magenta) and GFP-Cdc42EP5 (green). Cell was seeded on glass and imaged every 1 s (60s total). Right panels show merged and single channel magnifications (greyscale) of peripheral and perinuclear and peripheral regions.

**Movie EV4. Cdc42EP5 regulates focal adhesion dynamics.** Time lapse movies of 690.cl2 cells expressing GFP-Paxillin (grey) after transfection with control (siCtr) or Cdc42EP5 (siEP5) siRNA. Cell was seeded on glass and imaged every 30 s (10 min total). Lower panels represent motion analysis images of the same areas generated by overlaying blue, green and red images from different time points.

