## Supplementary Information for "Cdc42EP5/BORG3 modulates SEPT9 to promote actomyosin function and melanoma invasion and metastasis"

**SUPPLEMENTARY INFORMATION FILE**

This file includes:

6 Supplementary Figures (EV1-6)
3 Supplementary Tables

In addition, description for the Supplementary Movies 1-4 can be found at the end of the document

Farrugia et al. Expanded View.  Figure EV1

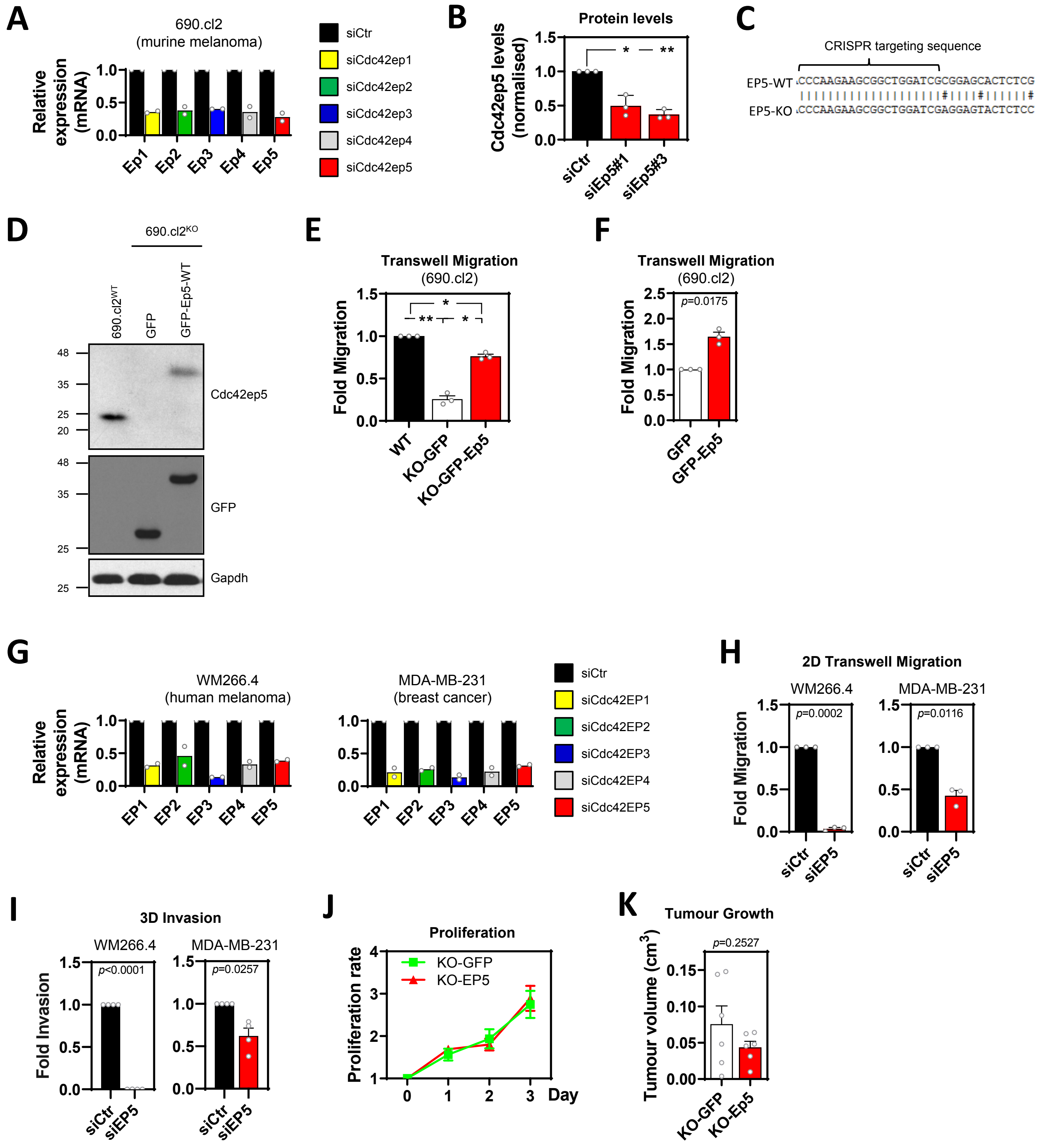

Farrugia et al. Expanded View.  Figure EV2

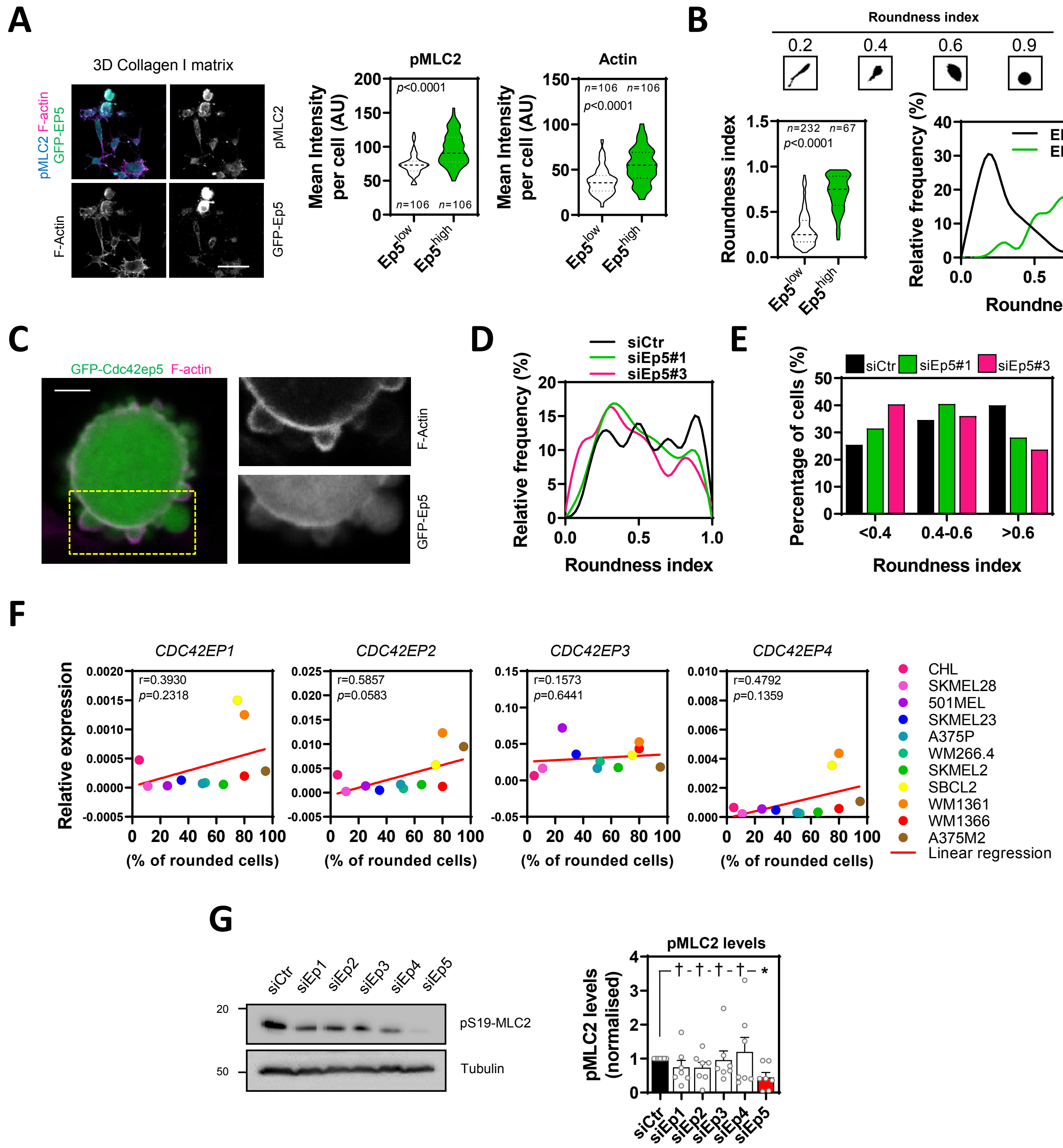

Farrugia et al. Expanded View. Figure EV3

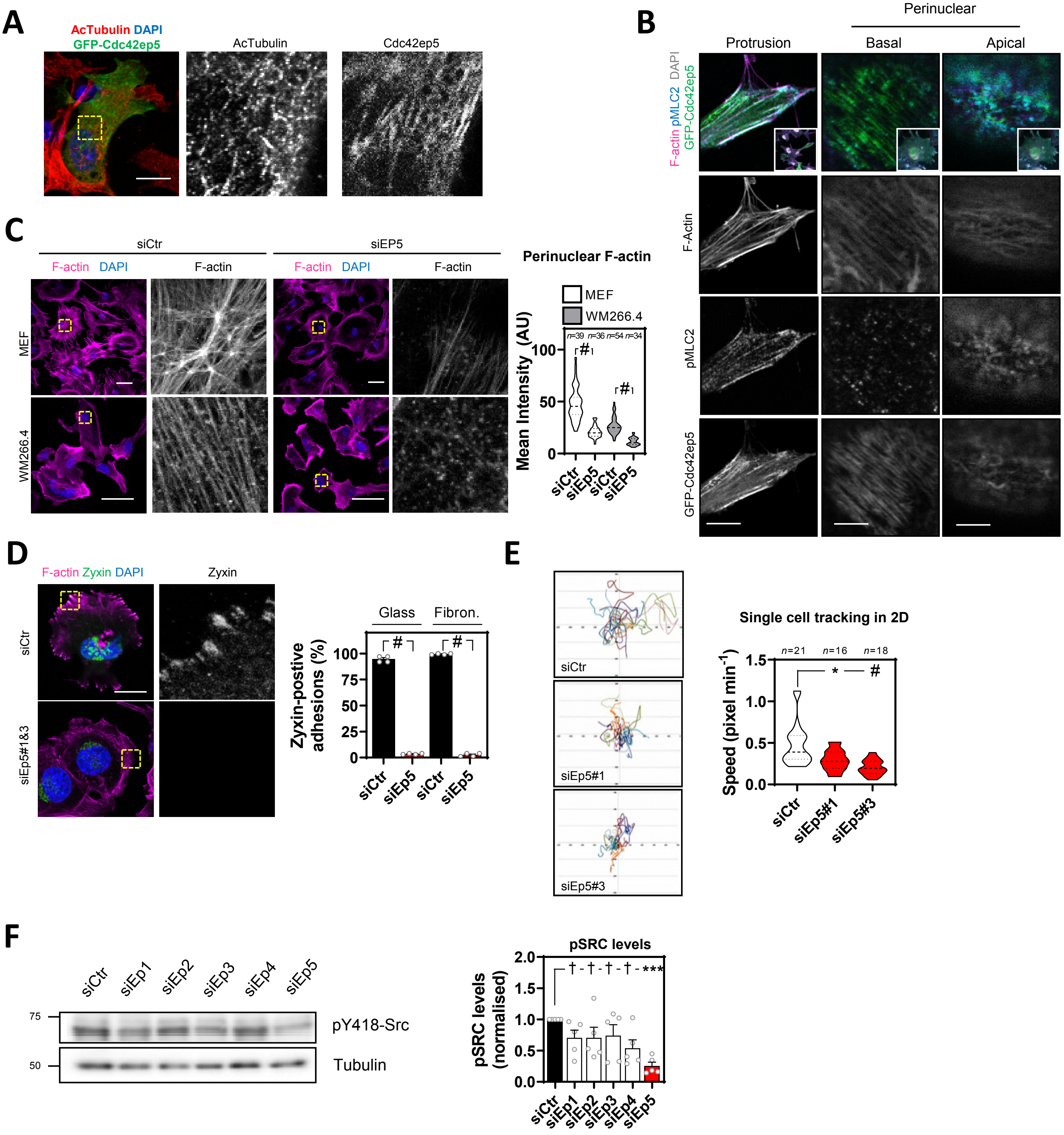

Farrugia et al. Expanded View. Figure EV4

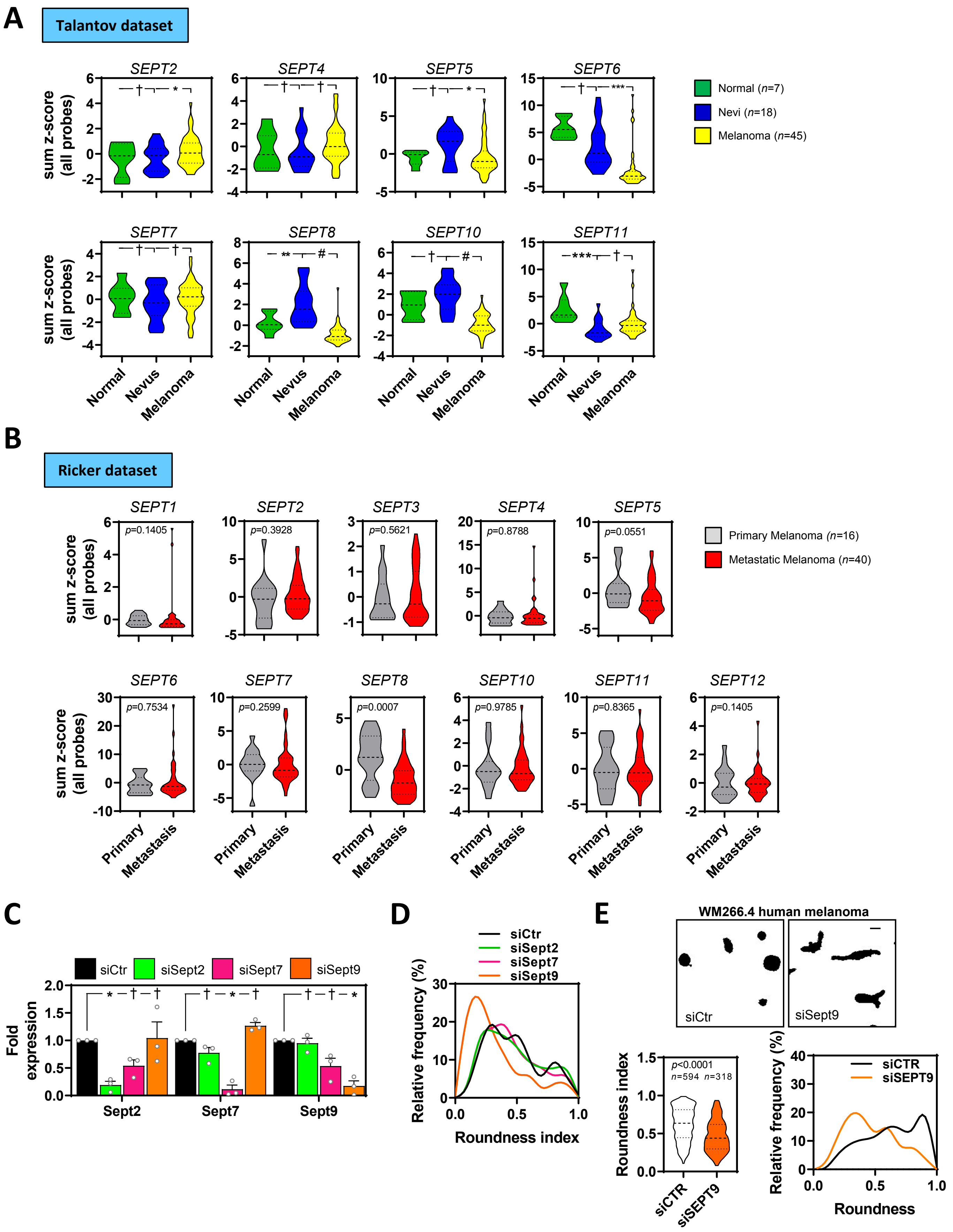

Farrugia et al. Expanded View. Figure EV5

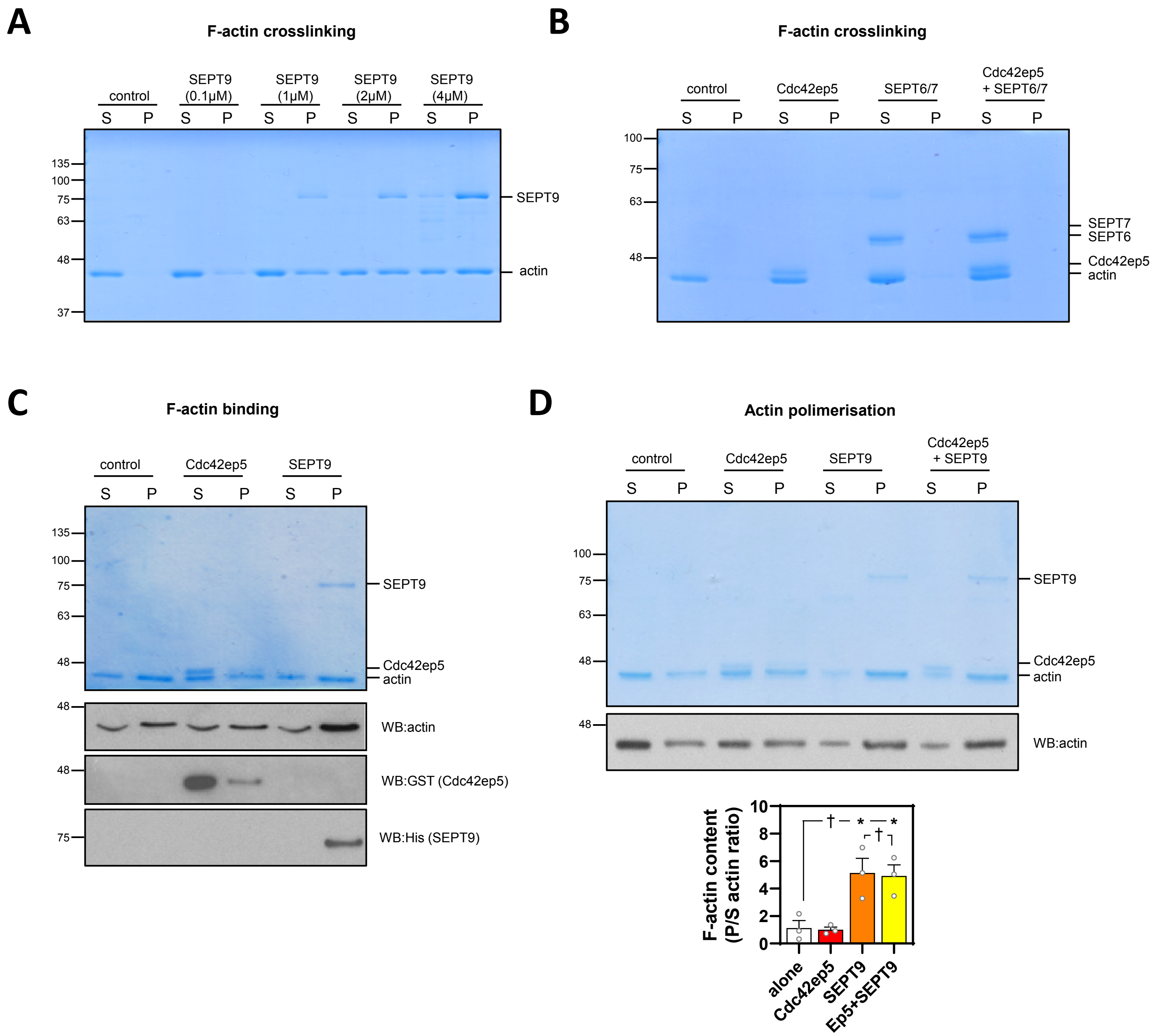

Farrugia et al. Expanded View. Figure EV6

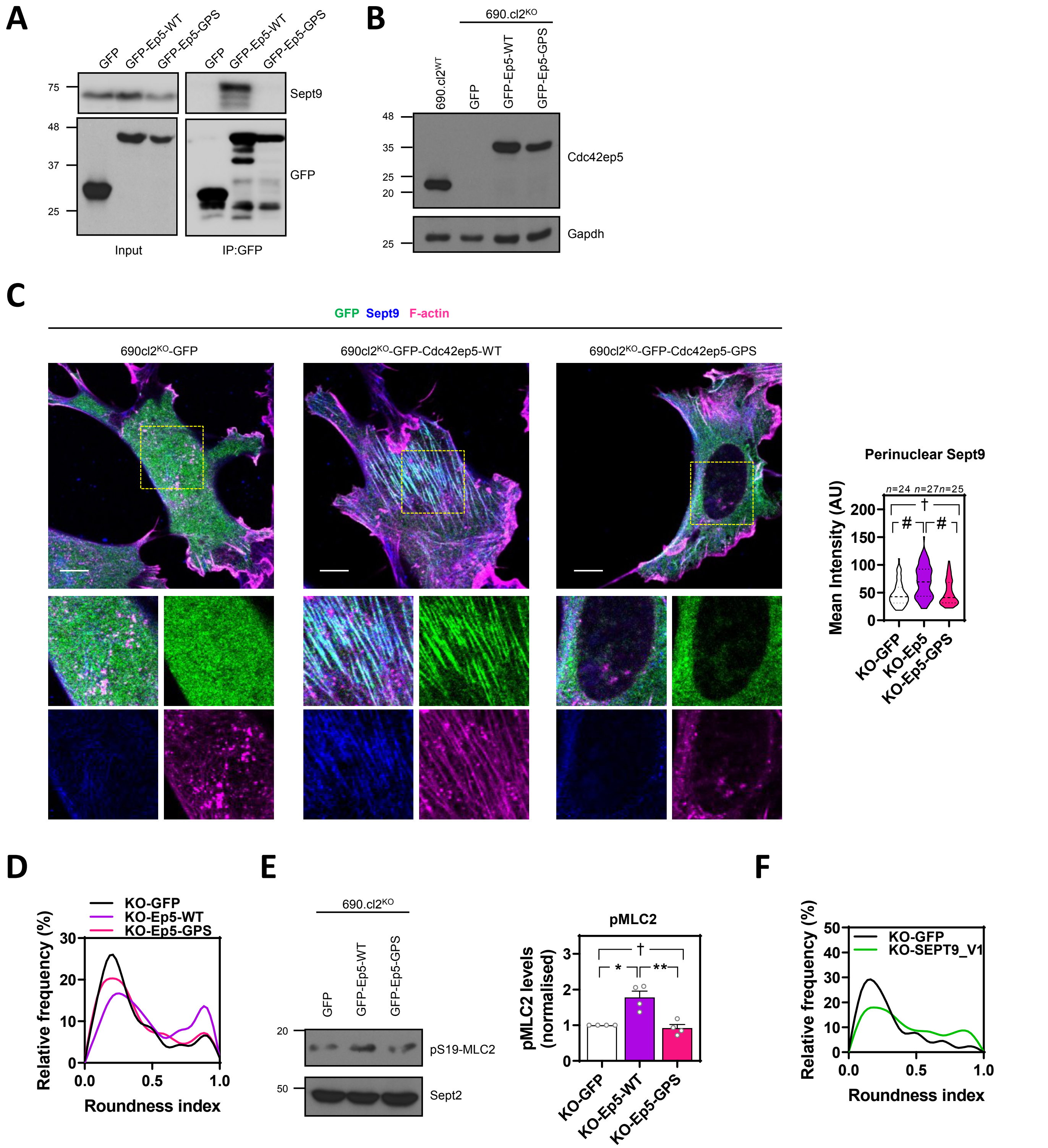

### Farrugia et al. Supplementary Information. Supplementary Table 1.

| Gene Name | Species | Target Sequence | Cat. Number |
|---|---|---|---|
| Cdc42ep1#1 | murine | GGAAAGAGACGCCUGACAG | D-045814-01 |
| Cdc42ep1#2 | murine | GGAGGCCGCUCUGGAAACA | D-045814-02 |
| Cdc42ep1#3 | murine | GCGAUAACCACUCUAGAUG | D-045814-03 |
| Cdc42ep1#4 | murine | CCGGCUAGCUCCACAGAUG | D-045814-04 |
| Cdc42ep2#1 | murine | GACCUUCCCUUCCAGUUUA | D-044823-01 |
| Cdc42ep2#2 | murine | GAUUAUGGAUCACGACCUA | D-044823-02 |
| Cdc42ep2#3 | murine | UGGCGGAGAUGACAUGUUU | D-044823-03 |
| Cdc42ep2#4 | murine | CGUGCAGAUUCCUACAUA | D-044823-04 |
| Cdc42ep3#1 | murine | GGAGCAAAGUAGUCUAUUA | D-046421-01 |
| Cdc42ep3#2 | murine | GAAAGCGGCUAACAACAAG | D-046421-02 |
| Cdc42ep3#3 | murine | GAUCUUGGGCCUUCACUUU | D-046421-03 |
| Cdc42ep3#4 | murine | GGAUAUUUCCUUUCUUCAA | D-046421-04 |
| Cdc42ep4#1 | murine | GAAGAAGGCAGACGCUAGA | D-049988-01 |
| Cdc42ep4#2 | murine | GGACACGUCUUUCCUCACU | D-049988-02 |
| Cdc42ep4#3 | murine | GAAUUCUGCUUCAUGGAUG | D-049988-03 |
| Cdc42ep4#4 | murine | CGGAGMGGACUCGAGCAA | D-049988-04 |
| Cdc42ep5#1 | murine | GGAGCACUCUCGAUCUCAG | D-063228-01 |
| Cdc42ep5#2 | murine | GCUCAAAUGCUGACCUCCA | D-063228-02 |
| Cdc42ep5#3 | murine | CGACGUCACGGGUCUGUAG | D-063228-03 |
| Cdc42ep5#4 | murine | GGGAUGCCCACCCUAGAGU | D-063228-04 |
| CDC42EP1#1 | human | GAAAGAGGCGGCUGACUGC | D-017551-01 |
| CDC42EP1#2 | human | AUGAUGAGGUCAAGGUGUG | D-017551-02 |
| CDC42EP1#3 | human | GGAAAAGCCGCAUGACCGA | D-017551-03 |
| CDC42EP1#4 | human | GCCCAGUGGCUGAGGUGAA | D-017551-04 |
| CDC42EP2#1 | human | GCAGUGACAUGUUUGGCGA | D-012215-01 |
| CDC42EP2#2 | human | GCAUGCAGAUCCCCACAUA | D-012215-02 |
| CDC42EP2#3 | human | CGCCACACCAUUCAUAUUG | D-012215-03 |
| CDC42EP2#4 | human | CCAUCUAUCUGAAGCGUGG | D-012215-04 |
| CDC42EP3#1 | human | GAUGAGGUGCUGAAUGUAA | D-017358-01 |
| CDC42EP3#2 | human | CCAAUAACAAGAAAGGAAA | D-017358-02 |
| CDC42EP3#3 | human | GCUCUCAUGUUGCCCUUAU | D-017358-03 |
| CDC42EP3#4 | human | CGAUGUCUUUGGAGAUAUU | D-017358-04 |
| CDC42EP4#1 | human | CCAGUAAGCUGCCCAAGAG | D-013494-01 |
| CDC42EP4#2 | human | CAAGACAGCCAGACAAGGA | D-013494-02 |
| CDC42EP4#3 | human | GACACCUCCUUCCUCAAUA | D-013494-03 |
| CDC42EP4#4 | human | CCCAUGCUCUGGAGGAUGA | D-013494-04 |
| CDC42EP5#1 | human | CCACUUCUGUAUACAUAAA | D-017378-13 |
| CDC42EP5#2 | human | CUGUAUACAUAAACGGCCA | D-017378-14 |
| CDC42EP5#3 | human | ACAUAAACGGCCAAGGUGU | D-017378-15 |
| CDC42EP5#4 | human | CAAGGUGUGUGCCCGGAAA | D-017378-16 |
| Sept2#1 | murine | GCAGGAAAGUAGAGAAUGA | D-051534-01 |
| Sept2#2 | murine | GAUAGAGGCUUCGACUGUU | D-051534-02 |
| Sept2#3 | murine | UGAAGAGCAUAGCAUUAAA | D-051534-03 |
| Sept2#4 | murine | GCGAUACACAAUAAGGUGA | D-051534-04 |
| Sept7#1 | murine | CAAAUCAAGUGUACAGAAA | D-042160-01 |
| Sept7#2 | murine | CCAAUAACGUACACUAUGA | D-042160-02 |
| Sept7#3 | murine | CGACAUUAAUCAACUCAUU | D-042160-03 |
| Sept7#4 | murine | CAUUGGACAUUGAGUUUAU | D-042160-04 |
| Sept9#1 | murine | GCAAAGUGGUGAACAUUGU | D-048947-01 |
| Sept9#2 | murine | CCAACGGCAUUGACGUGUA | D-048947-02 |
| Sept9#3 | murine | CGAAGCCUACCGAGUGAAA | D-048947-03 |
| Sept9#4 | murine | CUAUCGAAGUUGAGAAUAC | D-048947-04 |

**Supplementary Table 1. siRNAs:** Name (target), species, sequence and catalogue number of the single siRNAs used in the study.

| Target | Sequence |
|---|---|
| mCdc42ep1_F | ATGTCTTCGGAGATACGTCCTT |
| mCdc42ep1_R | GACTCTGCGAACCTGTTGGA |
| mCdc42ep2_F | TCCCCATCTATTTGAAACGTGG |
| mCdc42ep2_R | CCGCTGTTCCTGGAAGGAG |
| mCdc42ep3_F | CCAAGACCCCAATTTACCTGAAA |
| mCdc42ep3_R | CCCTCTTTGCCGATGTGTATAGT |
| mCdc42ep4_F | CAGCTCTGTGAACTCGAAGC |
| mCdc42ep4_R | GCTAGTGAGGAAAGACGTGTCC |
| mCdc42ep5_F | GGGATGCCCACCCTAGAGT |
| mCdc42ep5_R | TGGAGGTCAGCATTTGAGCAG |
| hCDC42EP1_F | GGCCACCACTACCCAGAGAT |
| hCDC42EP1_R | CAGCATCCGCAAATTCAAAGG |
| hCDC42EP2_F | TCCACCAAGGTGCCCATCTAT |
| hCDC42EP2_R | TCCACCGATAACCGGGAGG |
| hCDC42EP3_F | CCTGAGGTTACGGCCAAGT |
| hCDC42EP3_R | CTGGGATCCTCAGCGGC |
| hCDC42EP4_F | GCGAGTCCTTGGACGAACAG |
| hCDC42EP4_R | GCAGGGACATGGCATTCTTG |
| hCDC42EP5_F | CGATCAGGCCGCTTCTT |
| hCDC42EP5_R | AGGGGTCCGGGCTAGAG |
| mGapdh_F | GTGCAGTGCCAGCCTCGTCC |
| mGapdh_R | GCCACTGCAAATGGCAGCCC |
| mRplp1-F | ACCGTGCCGGCAGTCTACAG |
| mRplp1-R | ATGTTGACATTGGCCAGAGCCTTG |
| hGAPDH_F | GGCAAATTCCATGGCACCG |
| hGAPDH_R | GCATCGCCCCACTTGATTTT |

**Supplementary Table 2. Primers**. Names and sequences (forward and reverse) of the paired oligos used in this study for qRT-PCR. The name contains the target gene; h stands for human, m for murine; F for forward and R for Reverse.

**Farrugia et al. Supplementary Information. Supplementary Table 3.**

| Target | Antibody | Company (Cat. No) | Dilution | |
|---|---|---|---|---|
| | | | W.B. | I.F. |
| Actin | Anti-Actin | Sima (A3853( | 1:5000 | |
| Acetylated alpha Tubulin | Anti-acTub [6-11B-1] | Abcam (ab24610) | | 1:100 |
| Cdc42EP5 | Anti-Cdc42EP5 | Invitrogen (PA5-39085) | 1:500 | |
| Myosin light chain 2 | Anti-MLC | Cell Signalling (3672) | 1:2000 | |
| GAPDH | Anti-GAPDH (14C10) | Cell Signalling (3683) | 1:2000 | |
| GST | Anti-GST (91G1) | Cell Signalling (2625) | 1:2000 | |
| His probe | Anti-His (H3) | Santa Cruz (sc8036) | 1:1000 | |
| Phospho-focal adhesion kinase (Tyr397) | Anti-FAK (pY397) | Cell Signalling (8556) | 1:1000 | |
| Phospho-myosin Light Chain 2 (Ser19) | Anti-MLC2 pS19 | Cell Signalling (3671) | 1:500 | 1:100 |
| Phospho-Src (Tyr418) | Anti-Src p(Y418) | Invitrogen (44660G) | 1:1000 | |
| Phospho-paxillin (Tyr118) | Anti-paxillin pY118 | Life Technologies (44-722g) | 1:1000 | 1:100 |
| Septin 2 | Anti-SEPT2 | Proteintech (60075-1-Ig) | 1:1500 | |
| Septin 7 | Anti-SEPT7 | Proteintech (13818-1-AP) | 1:1500 | |
| Septin 9 | Anti-SEPT9 | Proteintech (10769-1-AP) | 1:1500 | 1:100 |
| Total Paxillin | Anti-Paxillin | BD Transduction Labs (610051) | 1:1000 | |
| Tubulin | Anti-β - Tubulin 1 | Sigma (T7816) | 1:20000 | |
| Zyxin | Anti-ZYX | Abcam (ab71842) | | 1:100 |

**Supplementary Table 3. Antibodies**. Name, company, catalogue number and working dilutions of all the antibodies used in the study.

**Supplementary Movie 1. Intravital imaging of 690.cl2 cells expressing Cdc42EP5 in a living tumour.** 690.cl2-KO cells ectopically expressing GFP-Cdc42EP5 (green) were grown subcutaneously and imaged in vivo using two photon microscopy, which enable the acquisition of second harmonic signal (collagen fibres, magenta). Images were acquired every 14 min for a total of 2h 45min. Scale bar, Scale bar, 100 µm. Middle panels show magnifications of indicated areas showing fast amoeboid movement into the matrix. Right panels represent motion analysis images of the same area generated by overlaying blue, green and red images from different time points. Distinct areas of colour indicate motile cells appear, whereas static regions appear white. .

**Supplementary Movie 2. Intravital imaging of 690.cl2 lacking Cdc42EP5 in a living tumour.** 690.cl2-KO cells ectopically expressing GFP (green) were grown subcutaneously and imaged in vivo using two photon microscopy, which enable the acquisition of second harmonic signal (collagen fibres, magenta). Images were acquired every 14 min for a total of 2h 45min. Scale bar, 100 µm. Middle panels show magnifications of indicated areas showing slow protrusive movement of cells and elongated morphology. Right panels represent motion analysis images of the same area generated by overlaying blue, green and red images from different time points. Distinct areas of colour indicate motile cells appear, whereas static regions appear white

**Supplementary Movie 3. Dynamic association between F-actin and Cdc42EP5.** Time-lapse movie of 690.cl2 cell expressing LIFEACT-RFP (magenta) and GFP-Cdc42EP5 (green). Cell was seeded on glass and imaged every 1 s (60s total). Right panels show merged and single channel magnifications (greyscale) of peripheral and perinuclear and peripheral regions.

**Supplementary Movie 4. Cdc42EP5 regulates focal adhesion dynamics.** Time lapse movies of 690.cl2 cells expressing GFP-Paxillin (grey) after transfection with control (siCtr) or Cdc42EP5 (siEP5) siRNA. Cell was seeded on glass and imaged every 30 s (10 min total). Lower panels represent motion analysis images of the same areas generated by overlaying blue, green and red images from different time points.
